# Combining genetic resources and elite material populations to improve the accuracy of genomic prediction in apple

**DOI:** 10.1101/2021.08.27.457920

**Authors:** Xabi Cazenave, Bernard Petit, François Laurens, Charles-Eric Durel, Hélène Muranty

**Author notes:** Corresponding author : Hélène Muranty.

## Abstract

Genomic selection is an attractive strategy for apple breeding that could reduce the length of breeding cycles. A possible limitation to the practical implementation of this approach lies in the creation of a training set large and diverse enough to ensure accurate predictions. In this study, we investigated the potential of combining two available populations, i.e. genetic resources and elite material, in order to obtain a large training set with a high genetic diversity. We compared the predictive ability of genomic predictions within-population, across-population or when combining both populations, and tested a model accounting for population-specific marker effects in this last case. The obtained predictive abilities were moderate to high according to the studied trait and were always highest when the two populations were combined into a unique training set. We also investigated the potential of such a training set to predict hybrids resulting from crosses between the two populations, with a focus on the method to design the training set and the best proportion of each population to optimize predictions. The measured predictive abilities were very similar for all the proportions, except for the extreme cases where only one of the two populations was used in the training set, in which case predictive abilities could be lower than when using both populations. Using an optimization algorithm to choose the genotypes in the training set also led to higher predictive abilities than when the genotypes were chosen at random. Our results provide guidelines to initiate breeding programs that use genomic selection when the implementation of the training set is a limitation.

## INTRODUCTION

Breeding programs in outbred fruit tree crops can take many years before new varieties are released, in part because of the long juvenile phase of the trees. Shortening the breeding cycle length for these crops could help increase genetic gain (van Nocker and Gardiner 2014). The length of the breeding cycle is also a constraint when breeders aim to introgress new traits from distant relatives or genetic resources, because several generations are usually needed before genotypes with the desired traits can be released as varieties.

In both cases, the use of molecular markers is an attractive strategy for the early identification of the most promising selection candidates or genetic resources (Myles 2013). Since most apple breeding programs around the world rely on a limited number of genotypes that are frequently used as parents for variety development (Noiton and Alspach 1996), mating strategies based on molecular markers could be used to broaden the genetic base of elite material (Yu *et al*. 2016) by identifying without phenotyping promising genetic resources that could be used as novel parents in pre-breeding programs (Crossa *et al*. 2016).

Recently, an approach referred to as genomic selection has particularly gained popularity among plant breeders (Voss-Fels *et al*. 2019), as genetic gain is expected to be higher with genomic selection compared to marker-assisted breeding, especially for complex traits (Xu *et al*. 2020). Genomic selection uses a training population of individuals that have been genotyped and phenotyped in order to estimate marker effects, then allowing the estimation of genomic breeding values of a candidate population that has only been genotyped (Meuwissen *et al*. 2001). The success of genomic selection depends on the accuracy of the predicted breeding values. The size of the training set (Zhang *et al*. 2017; Edwards *et al*. 2019), the relatedness between the training set and the candidates (Clark *et al*. 2012; Lehermeier *et al*. 2014) or the genetic architecture of the trait (Daetwyler *et al*. 2010; Wimmer *et al*. 2013) have been reported as factors affecting prediction accuracy. In fruit tree crops, the potential of genomic selection has been outlined (Kumar *et al*. 2012a; Nsibi *et al*. 2020) and its implementation could help in efficiently using genetic resources in apple (Kumar *et al*. 2020). However, initiating pre-breeding programs based on genomic predictions can be challenging, in part because maintaining and phenotyping populations of large size is time-consuming and costly. The establishment of a large training set could thus limit the implementation of genomic selection. A way to overcome this limitation could be to combine data coming from several breeding populations, like historical data or data from other countries. This strategy has been explored in animal breeding (Hayes *et al*. 2009; Ibánẽz-Escriche *et al*. 2009; Wientjes *et al*. 2015) and more recently in plant breeding (Technow and Totir 2015; Sverrisdóttir *et al*. 2018; Olatoye *et al*. 2020) generally showing little to no gain in prediction accuracy. This observation may result from differences in marker effects between the combined populations, as well as from the differences in relatedness between these populations and the candidates to predict. Prediction models that allow the estimation of population-specific marker effects have been proposed (Schulz-Streeck *et al*. 2012; Karoui *et al*. 2012; Lehermeier *et al*. 2015) but few studies have used such models in plant breeding (Rio *et al*. 2019; Ramstein and Casler 2019). Differences in LD patterns between the combined populations may also explain the low observed gains from combination. In this case, it has been suggested to use high-density marker data in order to ensure consistency in SNP-QTL linkage disequilibrium between populations (de Roos *et al*. 2008).

Multi-population training sets can also be interesting when the genotypes to predict result from crosses between the combined populations. If no phenotypic records are available for the progenies of such crosses (as can be the case when initiating a new breeding program or when crossing selection material with exotic germplasm), one could consider combining populations in order to increase the diversity of the training set (Brandariz and Bernardo 2019). In this context, an optimum proportion of each combined population may exist. If this occurs and the sizes of the populations differ, it can be relevant to use a subset of the populations rather than to combine all the genotypes into a unique training set. When the size of the training set has to be fixed in advance, different algorithms exist to optimize the composition of the training set (Rincent *et al*. 2012; Akdemir *et al*. 2015; Mangin *et al*. 2019; Ou and Liao 2019) and they could be used to choose the genotypes to include in a training set in each of the combined populations (Isidro *et al*. 2015).

In this study, we considered the two abovementioned scenarios to investigate the interest of combining different populations into a unique training set for genomic prediction in apple. First, we assessed the potential increase in prediction accuracy when combining two populations, namely genetic resources and elite material, instead of using only one of the two, either using a standard GBLUP model or a model that allows the marker effects to differ between populations. We then predicted the GEBV of the progenies from crosses between elite material and genetic resources and evaluated the prediction accuracy when using different proportions of genetic resources and elite material in the training set and different strategies to choose the genotypes of this training set.

The objectives of our study were to (1) compare the predictive ability of genomic prediction models using simple or combined training sets within and across populations, (2) investigate the effect of different proportions of the two populations used in a training set and (3) assess the impact of high and medium marker density in these cases.

## MATERIAL AND METHODS

In this study, three datasets were used for genomic predictions: the first one (hereafter referred to as the FBo-Hi dataset) regroups data coming from two past European projects while the second dataset (from here on referred to as the REFPOP dataset) contains data from an ongoing European project. The genotypes in both the FBo-Hi and REFPOP datasets are either old varieties (called genetic resources from now on) or progenies from breeding programs (called elite material in this study). Due to the different experimental design between the panels of the two datasets (see below), the FBo-Hi and REFPOP datasets provide an opportunity to investigate the effect of genomic predictions with or without the presence of genotype by environment interactions.

The third dataset contains data from crosses between old and modern varieties, which are part of a pre-breeding initiative engaged at IRHS, INRAE Angers. In this study, this dataset was only used as a validation set.

### Plant material

#### FBo-Hi panel

The panel regroups two different apple populations, from here on referred to as the genetic resources and elite material of the FBo-Hi dataset, for which phenotypic and genotypic data are available. The genetic resources represent European apple germplasm preserved in six European core-collections along with data coming from the EU-FP7 FruitBreedomics (FBo) project described in Laurens et *al*. (2018), while the elite material consists of progenies originating from biparental combinations from six European breeding programs that were gathered for pedigree-based QTL analysis during the HiDRAS (Hi) project (Kouassi *et al*. 2009). A total of 1194 unique genotypes (mainly old dessert apple cultivars) from the genetic resources were genotyped and phenotyped for at least one trait (see below). Similarly, 1018 progenies from 23 biparental combinations were genotyped and phenotyped for at least one trait. The genotypes of the FBo-Hi panel were not replicated across sites, except for some genotypes that were used to adjust phenotypic data.

#### REFPOP panel

The apple REFPOP is described in detail in Jung *et al*. (2020). This panel consists of 269 cultivars (hereafter referred to as genetic resources of the REFPOP) that are representative of the worldwide apple genetic diversity and of 265 progenies (hereafter referred to as elite material of the REFPOP) originating from 27 biparental combinations from various European breeding programs. Some of the genotypes of the REFPOP panel are also part of the FBo-Hi panel, as 189 accessions and 155 progenies are found in both panels. The panel is replicated across six locations in six European countries with contrasting environments and each genotype is replicated at least twice in each environment. At each site, the genetic resources and elite material are planted in the same orchard.

#### Panel of hybrids

The panel consists of 473 so-called ‘hybrids’ originating from 10 biparental combinations of approximately the same size. Each combination involved a controlled cross between an old cultivar and an elite cultivar. The mating design is presented in Table S1 and involved 5 old cultivars and 9 elite cultivars. Each hybrid genotype was represented by only one tree in the orchard in Angers, France.

### Genotypic data

#### Genotyping of the plant material

For both the FBo-Hi and REFPOP panels, the genetic resources were genotyped using the Affymetrix Axiom Apple 480K SNP genotyping array (Bianco *et al*. 2016) and the elite material using the Illumina Infinium 20K SNP genotyping array (Bianco *et al*. 2014). The hybrids were also genotyped with the 20K genotyping array. After filtering, 7,060 SNP markers were retained from the 20K array and 303,239 SNP markers from the 480K array. More information about the filtering and quality check procedure can be found in Jung *et al*. (2020).

#### Genotype imputation

For the elite and hybrids genotypes, the 20K SNPs were completed to reach the 480K genotyping array density by imputation with BEAGLE 4.0 software (Browning and Browning 2007), which can use a reference set of phased marker data along with pedigree information to improve the imputation quality. The reference panel proposed by Jung et al. (2020) was used in this study. Missing SNP marker data in the reference panel were first imputed and haplotypes were phased using the default parameters of BEAGLE 4.0. The phased marker data along with pedigree information inferred in Muranty et al. (2020) and updated in an ongoing apple pedigree project (Howard *et al*. 2018) were then provided to the software for the imputation from medium to high-density.

### Phenotypic data

#### FBo-Hi dataset

The genetic resources were phenotyped between 2012 and 2014 in six European research institutes. Several fruit quality and phenology traits were measured by assessor pairs according to the recommendations of the European Cooperative Programme for Plant Genetic Resources (Watkins *et al*. 1982) and the notation for each trait was harmonized between institutes. When available, each institute also provided FBo-Hi notations for the measured traits (Table S2).

The elite material was phenotyped between 2003 and 2005 in six European countries. Several traits linked to the productivity and fruit quality were measured. More details about the phenotyping procedure can be found in Muranty *et al*. (2015) and Kouassi *et al*. (2009).

Five traits were phenotyped in both populations: harvest date, fruit over-color, fruit juiciness, fruit acidity and fruit crispness. As we were interested in combining phenotypic information from both populations, only these traits were used for the genomic predictions. When the traits in the two populations were evaluated using different ordinal scales, a correspondence table between the two scales was created in order to use phenotypic data that could be combined.

#### REFPOP dataset

Traits related to yield, fruit quality and phenology were measured between 2018 and 2020 using a common protocol in each location. The detailed protocol is described in Jung *et al*. (submitted). Harvest date and fruit over-color were the only two traits that were measured in both the FBo-Hi and the REFPOP panels.

#### Dataset of hybrids

The hybrids were phenotyped in 2019 and 2020. Fruit weight, fruit number, fruit over-color and harvest date were measured for each tree following the protocol proposed for the REFPOP dataset.

#### Data adjustment

The phenotypic data of the REFPOP dataset and the genetic resources of the FBo-Hi dataset were adjusted for year and site effects and those of the hybrids were adjusted for year effect, as they were evaluated in only one site. To do so, the estimated marginal means of the raw phenotypic data were computed using the emmeans function of the emmeans package (Russel 2021). Prior to this adjustment, the phenotypic data of the REFPOP dataset were also corrected for spatial heterogeneity as in Jung et al. (2020) using the P-spline ANOVA approach with the PSANOVA function of the SpATS package (Rodríguez-Álvarez *et al*. 2018).

Heritability of genotypic means was estimated in both cases as 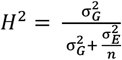, where 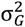 is the variance of genotype effects, 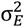 is the residual variance and *n* is the mean number of observations per genotype.

In the case of the elite material of the FBo-Hi dataset, the Best Linear Unbiased Predictors (BLUPs) of clonal values computed in Kouassi et al. (2009) were used as phenotypic values.

### Genetic characterization of the populations

#### Structure and genetic diversity across the populations

The structure of the populations was investigated through a principal component analysis (PCA) performed on the SNP marker data of the hybrids, the genetic resources and the elite material. A PCA was thus performed for the FBo-Hi dataset and another one for the REFPOP dataset. To reduce the computational time of the analysis, the marker data were first pruned based on linkage disequilibrium using the indep-pairwise function in the PLINK 1.9 program (Purcell *et al*. 2007), by pruning the markers with a pairwise r^2^ > 0.1 in a 50kb window. 12,363 SNP markers were retained after pruning from the FBo-Hi dataset and 12,290 markers from the REFPOP dataset. The PCA was carried out with the prcomp function from the R *stats* package. To further assess the differentiation between the elite material, the genetic resources and the hybrids, pairwise F_ST_ values between the populations were computed using the pairwise.WCfst function from the R *hierfstat* package (Goudet 2005).

The allele frequencies along the genome of the elite material and genetic resources were compared using sliding windows for both datasets. For each chromosome, windows of 2Mb with a shift of 400kb were built and the average minor allele frequency of the SNPs included in the windows were computed for the genetic resources. The frequency of the minor allele in the genetic resources was then computed in the elite material following the same procedure. When the number of SNPs of a window was less than 200, the mean minor allele frequency was set to missing.

#### Linkage disequilibrium

For each population of the FBo-Hi and REFPOP datasets, the linkage disequilibrium (r^2^) was computed as the square correlation coefficient between pairs of markers within a 500kb distance using the r^2^ function of the PLINK 1.9 program. Marker pairs were placed into bins of 500bp according to their pairwise distance and the LD decay was plotted as the mean r^2^ of each bin.

#### Evaluated scenarios for genomic predictions

We evaluated the interest of population combination for two purposes: in the first case, we predicted a population (genetic resources or elite material) using either the same population (within-population prediction), the other population (across-population prediction, AP) or a training set (TS for short) including genotypes from both populations (combination prediction, Comb). In the second case, we predicted the genetic resources x elite material hybrids previously described with a training set combining genotypes from the genetic resources and elite material populations (Prop_hybrids scenario). In the latter case, we investigated the effect of different proportions of the two combined populations on the predictive ability given different TS sizes. The explored scenarios are summarized in Table 1.

**Table 1.** Composition of the training and validation set in the different scenarios

| Method | Training set | Validation set |
| --- | --- | --- |
| WP | Genetic resources | Genetic resources |
|  | Elite | Elite |
| AP | Elite | Genetic resources |
|  | Genetic resources | Elite |
| Comb | Genetic resources + Elite | Genetic resources |
|  | Genetic resources + Elite | Elite |
| Prop_hybrids | Genetic resources + Elite | Hybrids |

#### Genomic prediction models

Two different prediction models were evaluated in this study. For both models, the Genomic Estimated Breeding Values (GEBV) of the candidates were calculated using the following mixed model:

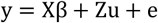

where y is a vector of phenotypic values, X is an incidence matrix relating fixed effects to the observations, β is the grand mean, Z is an incidence matrix linking the observations to the breeding values, u is a vector of breeding values for each individual and e is the vector of residuals.

#### Standard GBLUP model

We used the GBLUP model which is derived from the mixed model presented above with 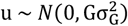 and 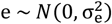. 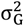 and 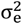 are respectively the genetic and residual variances and G is the Genomic Relationship Matrix (GRM, see below). The variance components and the breeding values were estimated with the *kin*.*blup* function of the R rrBLUP package (Endelman 2011).

#### Multi-group GBLUP model

A second model that takes the genomic correlation between the populations into account was used. In contrast to the standard GBLUP, this model allows the marker effects to be different (but correlated) between the two populations. Following Lehermeier *et al*. (2015),the model can be written as:

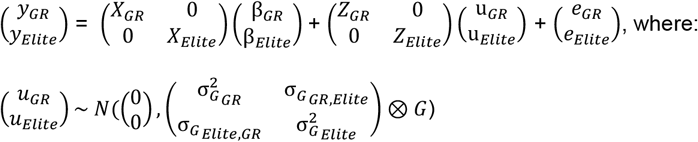

Here *X*_*GR*_ and *X*_*Elite*_ are incidence matrices relating fixed effects to the observations in each population, β_*GR*_ and β_*Elite*_ represent the mean of the two populations, *Z*_*GR*_ and *Z*_*Elite*_ are the incidence matrices relating the breeding values to the observations in each population, u_*GR*_ and u_*Elite*_ are the vectors of breeding values of the genetic resources and the elite material, 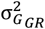 and 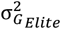 are the genomic variances of the genetic resources and the elite material, 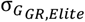 is the genomic covariance between the two populations and *e*_*GR*_ and *e*_*Elite*_ are vectors of residuals for each population. The genomic correlation between the two populations, defined as the correlation of the marker effects of each population, was computed from the estimated parameters as:

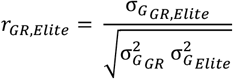

The genetic variances and covariances were estimated using a Gibbs sampler implemented in the MTM package. The Gibbs sampler was run for 20,000 iterations, the first 4000 being discarded as burn-in. One in every two samples was kept and the genetic parameters were then estimated by computing the posterior means of the remaining samples.

#### Genomic relationship matrix

For both models, the genomic relationship matrix (GRM) G was estimated as in VanRaden (2008):

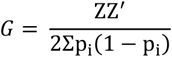

where Z = M – 2*p*_*i*_ is the centered matrix of the marker data, M is the matrix of marker data and p_i_ is the minor allele frequency at locus *i*. Here Z corresponds to the marker data of both genetic resources and elite material combined into one dataset, allowing the estimation of the relationships within and across populations. The genomic relationship matrix was calculated with the *A*.*mat* function of the rrBLUP package.

The GRM used in the models was computed according to the populations used in the training and validation sets as presented in Table 1. In the case of within-population prediction, the marker data of the predicted population was used to compute the GRM, while the marker data of the two populations was used to compute the GRM for the across-population prediction and combination prediction cases. To evaluate the influence of the marker density on the predictive ability of the models, the GRM was computed using the high-density marker data and a subset corresponding to the SNP markers of the 20K genotyping array in each case.

#### Predictive ability

The predictive ability of the different models was evaluated as the Pearson correlation coefficient between the GEBV and the phenotypic values of the individuals in the validation set. In a first step, the elite material and genetic resources were considered separately and a fivefold cross-validation scheme was used to randomly split the genotypes between training and validation sets within each population, allowing within-population genomic predictions (from here on referred to as WP method). The same validation sets (VS) were also predicted using all the genotypes in the opposite population (across-population prediction, from here on referred to as AP method) or by combining the genotypes of the two populations into a unique training set (combined-populations prediction, Comb and MG-Comb methods, see below). In this case, both the standard GBLUP (Comb method) and multi-group GBLUP (MG-Comb method) models described above were used. This procedure was replicated 20 times for both the FBo-Hi and REFPOP datasets. The approach is summarized in Figure 1A and the different TS and VS compositions are shown in Table 1.

**Figure 1.**
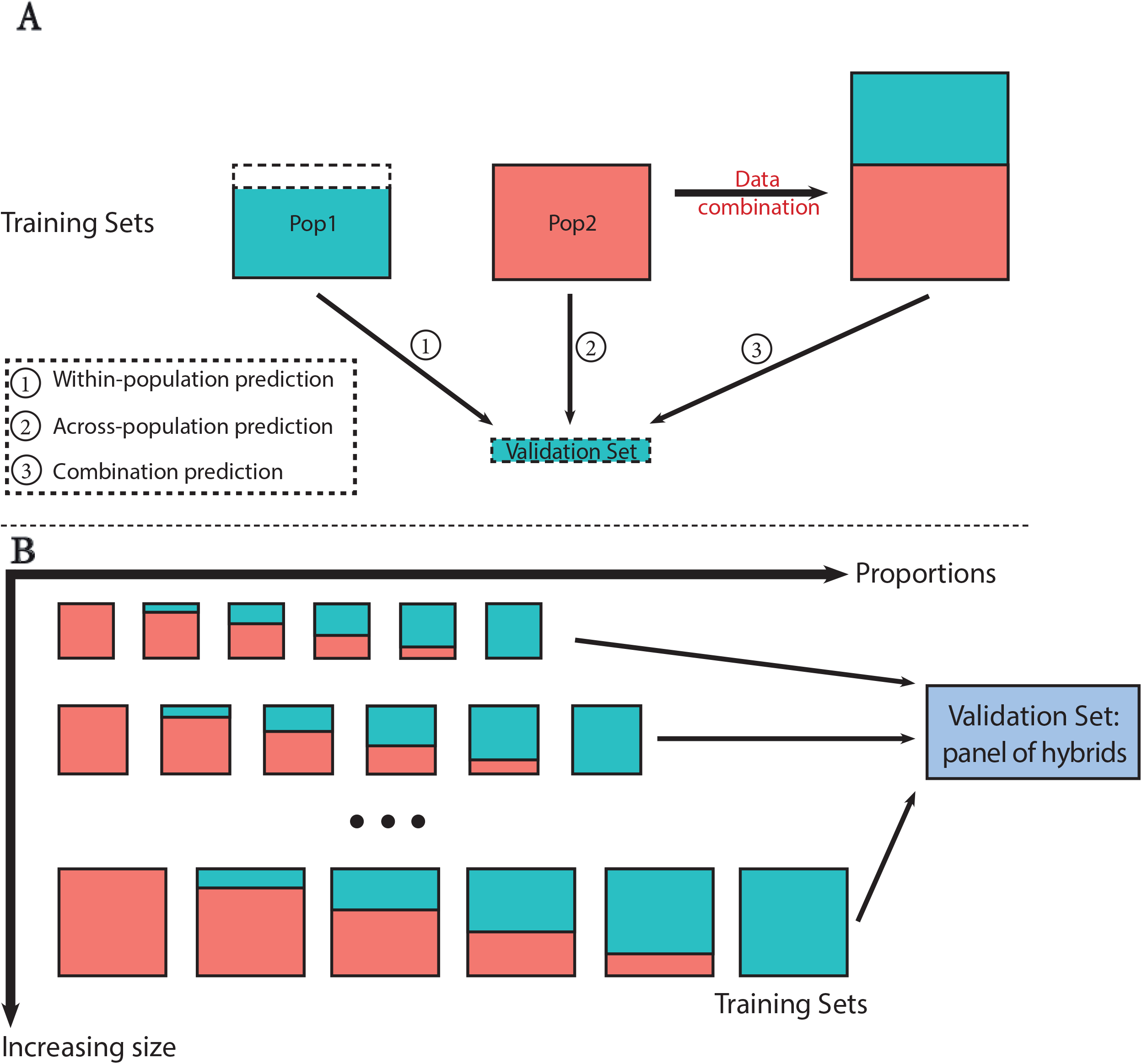
Schematic representation of the tested scenarios in this study. **A**. The validation set is composed of 20% of one of the two populations (elite material or genetic resources) and is predicted either by the remaining 80% of the population (within-population prediction), by the other population (across-population prediction) or by combining both populations (combination prediction). **B**. The panel of hybrids is used as validation set and is predicted by combining elite material and genetic resources into a unique population. Several training sets with increasing sizes and different proportions of the two populations are tested.

#### Proportion of GR/elite in the training set

When predicting the hybrids, the influence of the elite/GR proportion in the TS was also investigated (Prop_hybrids scenario in Table 1). To do so, the total size of the TS was first fixed and the TS was then built with increasing proportions of elite individuals, from 0% to 100% by steps of 20%, and correspondingly decreasing proportions of genetic resources individuals (Figure 1B). From here on, the tested proportions will be noted as “prop_elite_/prop_GR_ proportion”, where prop_elite_ and prop_GR_ are values between 0 and 1 that refer to the tested proportions (for example the 0.2/0.8 proportion means that the TS contains 20% of elite genotypes and 80% of genotypes from the genetic resources). For all the proportions, different TS sizes were evaluated: the TS size ranged from 50 to 500 by steps of 50 individuals when the genotypes in the TS were chosen from the FBo-Hi dataset and from 50 to 250 by steps of 50 individuals when using the REFPOP dataset. In this last case, the maximum TS size was smaller than for the FBo-Hi datasets because all the proportions could not be tested otherwise given the smaller number of genotypes in each population of the REFPOP. The individuals in the TS were chosen with 3 different methods, as described below. These methods were applied separately to the elite and GR populations by sampling *n*_*1*_ genotypes from the elite population and *n*_*2*_ genotypes from the genetic resources, where *n*_1_ and *n*_*2*_ depend on the studied proportion and *n*_*1*_ *+ n*_*2*_ *= n* is a predefined TS size as described above. The chosen individuals in the two populations were then combined into a unique TS that was used for the prediction of the hybrids The GEBV of the hybrids was predicted with the standard GBLUP model described above.

#### Method 1: choice of the TS based on the CDpop criterion

The CDpop criterion proposed by Rincent *et al*. (2017) was used to choose the individuals for the TS. CDpop is derived from the coefficient of determination (Rincent *et al*. 2012), which is a measure of the expected reliability of the predictions. The optimization based on CDpop aims to maximize the mean CD of the contrast matrix of the validation set to predict, as defined in Rincent *et al*. (2017). For the two populations, an initial number *n*_*1*_ (respectively *n*_*2*_) of genotypes was first sampled. An exchange algorithm was then used to replace an individual within each group by an individual from the same population that had not been included yet, and this latter was kept in the group if the mean CD of the group increased after its inclusion. With our data, we found that the mean CD reached a plateau after 1000 to 5000 iterations. Since the choice of the TS based on CDpop is based on the initial sampling, the final TS composition can vary. The method was thus repeated 20 times with 3000 iterations.

#### Method 2: choice of the TS based on relatedness

For each population, the genotypes were chosen based on their relationship with the validation set. To do so, we calculated the mean relationship coefficient between an individual and the candidates from the GRM and chose the genotypes with the highest mean relationship to be part of the TS.

#### Method 3: choice of the TS by random sampling

In the “Random_strat” method, the TS was constituted by combining *n*_*1*_ genotypes sampled from the elite population and *n*_*2*_ genotypes sampled from the genetic resources. The “Random” method is similar except that *n* genotypes were sampled to constitute the TS, regardless of the population to whom they belonged. The two methods were applied 100 times.

## RESULTS

### Characterization of the populations

In order to investigate the potential of combining genetic resources and elite material into a training set, we first investigated the genetic and phenotypic differences between the two populations. Since the results were very similar between the two datasets, we only present the results obtained with the FBo-Hi dataset. See Supplementary Data for the same analyses with the REFPOP dataset.

Figure 2 shows the first two principal components from the PCA obtained using the pruned SNP marker data. A clear distinction between the two populations can be observed, although some genetic resources (for example Golden Delicious, Jonathan or Cox’s Orange Pippin) appear to be closer from the elite material than from the other genetic resources. As expected, the hybrids were plotted between the two populations. The F_ST_ value computed for the marker data indicated a low differentiation between the genetic resources and elite material (F_ST_ = 0.023, see Table S3). The linkage disequilibrium decay was rapid in both genetic resources and elite material (Figure S1), the decay being faster in the genetic resources with an average r^2^ calculated at 1kb, 5kb, 100kb and 250 kb of 0.24, 0.2, 0.18 and 0.15 in the genetic resources and 0.29, 0.26, 0.22 and 0.2 in the elite material in the FBo-Hi dataset. Very similar values were observed in the REFPOP dataset. In the two panels, allelic frequencies were similar between elite material and genetic resources for the largest part of the genome (Figure S2-3), although some major differences could be observed for some genomic regions (for example in chromosome 3 and 15).

**Figure 2.**
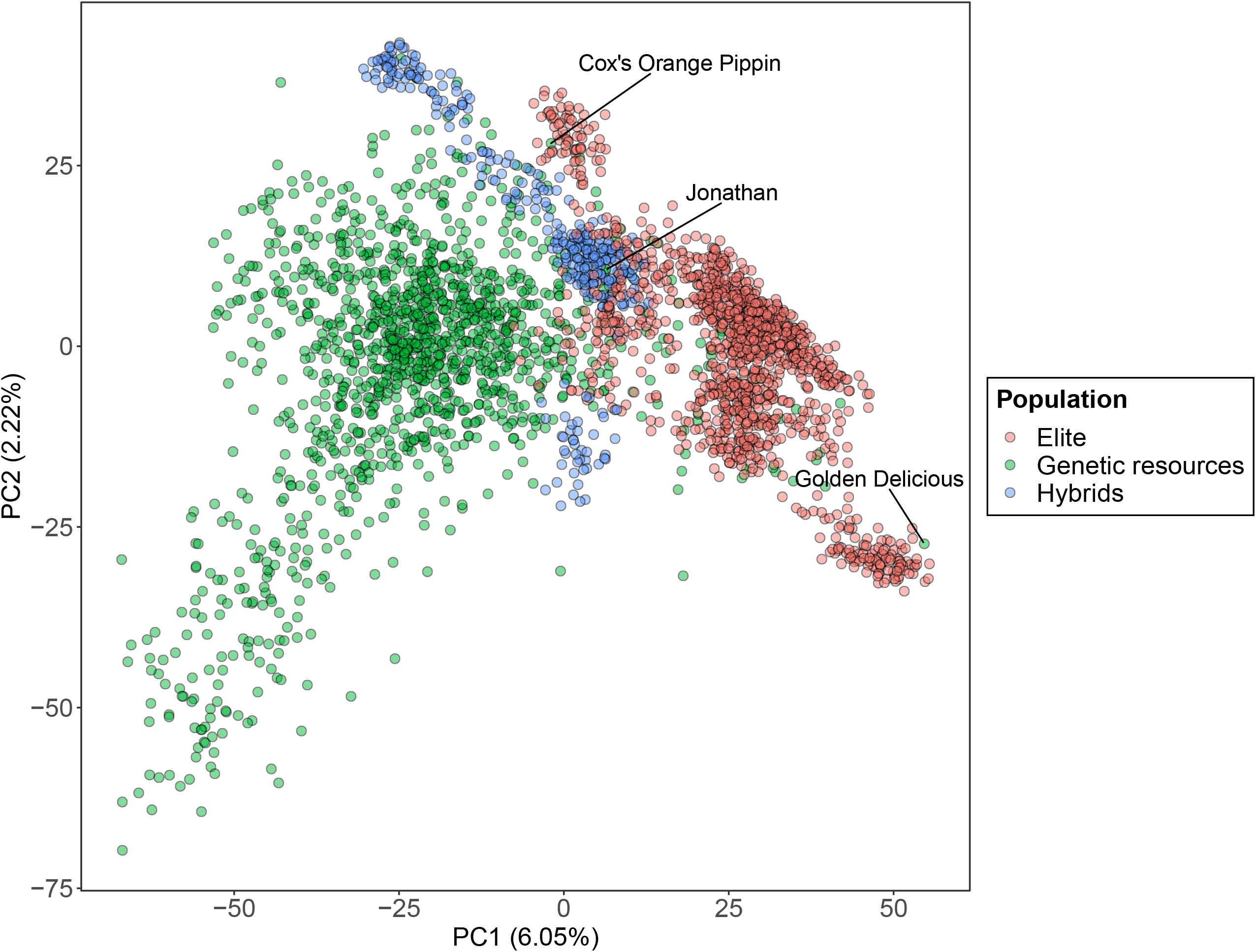
Principal Component Analysis (PCA) performed using pruned marker data of the FBo-Hi and hybrids panel.

Figures S4-6 show that the phenotypic distribution is different for some traits between the two populations. Fruits phenotyped in the elite material were juicier and crispier than in the genetic resources, and there were more fruits per tree in the elite material. The mean fruit over-color was also slightly higher in the elite material, although the same bimodal distribution could be observed in the two populations. The mean fruit acidity, fruit weight and harvest date were similar for both populations, but a wider range in acidity score was measured in the elite material compared to genetic resources, while the range in harvest date was larger in the genetic resources. The fruit weight distribution was extremely similar in both populations. In both datasets, the heritabilities were generally high (>0.7) for all the traits (Figure S7 and Table S2).

The genomic correlations between the two populations were moderate to high, ranging from 0.42 to 1 (Table 2). The lowest genomic correlations were measured for fruit weight, fruit crispness and fruit juiciness. The correlation was high for fruit over-color and acidity (around 0.7 in both cases) and was highest for harvest date (in both datasets) and fruit number, with a correlation of almost 1.

**Table 2.**
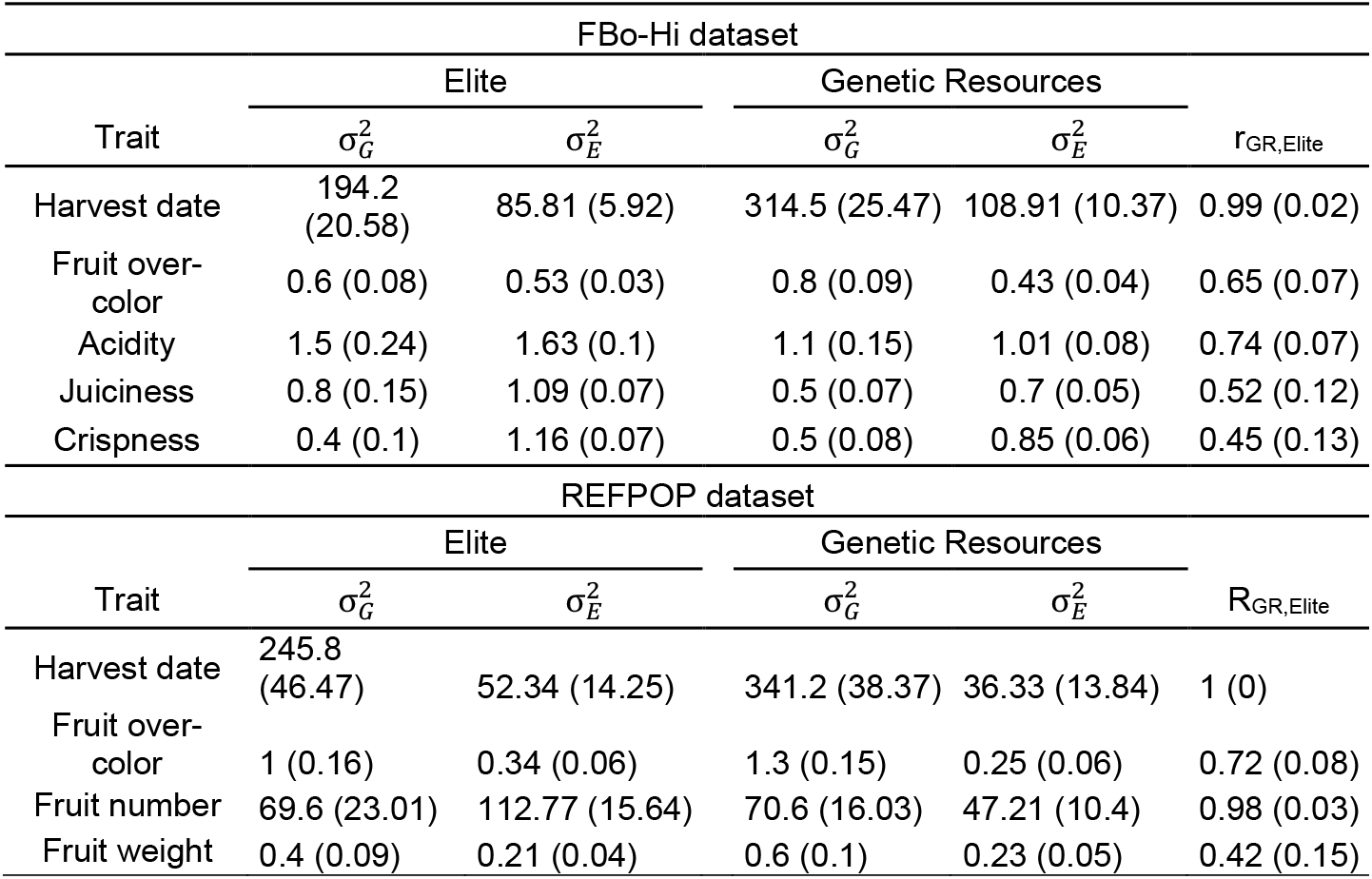
Genomic and environmental variances and genomic correlations estimated from the Gibbs sampler for each trait in the two population of the FBo-Hi and REFPOP datasets

| FBo-Hi dataset |  |  |  |  |  |
| --- | --- | --- | --- | --- | --- |
| Trait | Elite | | Genetic Resources | | $r_{GR,Elite}$ |
| | $\sigma_G^2$ | $\sigma_E^2$ | $\sigma_G^2$ | $\sigma_E^2$ | |
| Harvest date | 194.2<br>(20.58) | 85.81 (5.92) | 314.5 (25.47) | 108.91 (10.37) | 0.99 (0.02) |
| Fruit over-color | 0.6 (0.08) | 0.53 (0.03) | 0.8 (0.09) | 0.43 (0.04) | 0.65 (0.07) |
| Acidity | 1.5 (0.24) | 1.63 (0.1) | 1.1 (0.15) | 1.01 (0.08) | 0.74 (0.07) |
| Juiciness | 0.8 (0.15) | 1.09 (0.07) | 0.5 (0.07) | 0.7 (0.05) | 0.52 (0.12) |
| Crispness | 0.4 (0.1) | 1.16 (0.07) | 0.5 (0.08) | 0.85 (0.06) | 0.45 (0.13) |
| REFPOP dataset |  |  |  |  |  |
| Trait | Elite | | Genetic Resources | | $R_{GR,Elite}$ |
| | $\sigma_G^2$ | $\sigma_E^2$ | $\sigma_G^2$ | $\sigma_E^2$ | |
| Harvest date | 245.8<br>(46.47) | 52.34 (14.25) | 341.2 (38.37) | 36.33 (13.84) | 1 (0) |
| Fruit over-color | 1 (0.16) | 0.34 (0.06) | 1.3 (0.15) | 0.25 (0.06) | 0.72 (0.08) |
| Fruit number | 69.6 (23.01) | 112.77 (15.64) | 70.6 (16.03) | 47.21 (10.4) | 0.98 (0.03) |
| Fruit weight | 0.4 (0.09) | 0.21 (0.04) | 0.6 (0.1) | 0.23 (0.05) | 0.42 (0.15) |

### Predictive abilities when combining populations

Table 3 presents the predictive abilities for within and across-population predictions, as well as for predictions obtained when combining elite material and genetic resources into a unique training set for the FBo-Hi and REFPOP datasets.

**Table 3.** Predictive abilities of the measured traits in the within-population (WP), across-population (AP), combined populations (Comb) and MG-GBLUP methods

| Historical dataset |  |  |  |  |  |
| --- | --- | --- | --- | --- | --- |
| Trait | Method | Elite material |  | Genetic resources |  |
|  |  | Medium density | High density | Medium density | High density |
| Harvest date | WPP | **0.69 (0.12)** | 0.7 (0.12) | 0.80 (0.03) | 0.80 (0.04) |
|  | APP | 0.52 (0.12) | 0.54 (0.13) | 0.44 (0.07) | 0.49 (0.07) |
|  | Comb | **0.69 (0.12)** | **0.71 (0.11)** | **0.81 (0.03)** | **0.82 (0.03)** |
|  | MG-GBLUP | **0.69 (0.13)** | **0.71 (0.11)** | **0.81 (0.04)** | 0.81 (0.04) |
| Fruit over-color | WPP | 0.75 (0.06) | 0.75 (0.06) | 0.64 (0.07) | 0.61 (0.06) |
|  | APP | 0.60 (0.14) | 0.57 (0.14) | 0.48 (0.11) | 0.43 (0.12) |
|  | Comb | **0.77(0.06)** | **0.76 (0.06)** | **0.7 (0.08)** | **0.63 (0.07)** |
|  | MG-GBLUP | **0.77 (0.06)** | **0.76 (0.06)** | 0.67 (0.08) | **0.63 ((0.07)** |
| Crispness | WPP | 0.34 (0.06) | 0.33 (0.06) | 0.34 (0.05) | 0.34 (0.05) |
|  | APP | 0.15 (0.07) | 0.17 (0.06) | 0.21 (0.05) | 0.22 (0.05) |
|  | Comb | **0.36 (0.06)** | **0.36 (0.06)** | **0.35 (0.05)** | **0.36 (0.05)** |
|  | MG-GBLUP | 0.35 (0.06) | 0.35 (0.06) | **0.35 (0.04)** | 0.35 (0.05) |
| Juiciness | WPP | **0.49 (0.05)** | 0.48 (0.05) | **0.4 (0.05)** | 0.41 (0.05) |
|  | APP | 0.18 (0.07) | 0.16 (0.07) | 0.13 (0.07) | 0.14 (0.07) |
|  | Comb | **0.49 (0.05)** | **0.49(0.05)** | 0.38 (0.05) | 0.4 (0.05) |
|  | MG-GBLUP | **0.49 (0.05)** | **0.49 (0.05)** | **0.4 (0.05)** | **0.42 (0.05)** |
| Acidity | WPP | 0.48 (0.05) | 0.47 (0.05) | 0.45 (0.05) | 0.43 (0.05) |
|  | APP | 0.39 (0.05) | 0.39 (0.05) | 0.28 (0.06) | 0.23 (0.06) |
|  | Comb | **0.51 (0.04)** | **0.5 (0.04)** | **0.47 (0.06)** | **0.46 (0.06)** |
|  | MG-GBLUP | **0.51 (0.04)** | **0.49 (0.04)** | **0.47 (0.05)** | 0.45 (0.05) |
| REFPOP dataset |  |  |  |  |  |
| Trait | Method | Elite material |  | Genetic resources |  |
|  |  | Medium density | High density | Medium density | High density |
| Harvest date | WPP | **0.6 (0.09)** | 0.6 (0.09) | 0.79 (0.04) | 0.78 (0.04) |
|  | APP | 0.44 (0.12) | 0.45 (0.12) | 0.41 (0.08) | 0.47 (0.08) |
|  | Comb | **0.6 (0.10)** | **0.63 (0.10)** | **0.8 (0.04)** | **0.8 (0.04)** |
|  | MG-GBLUP | **0.6 (0.10)** | 0.62 (0.10) | 0.79 (0.04) | 0.79 (0.04) |
| Fruit over-color | WPP | 0.79 (0.05) | 0.78 (0.05) | 0.69 (0.06) | 0.64 (0.06) |
|  | APP | 0.73 (0.05) | 0.69 (0.06) | 0.56 (0.09) | 0.52 (0.09) |
|  | Comb | 0.81 (0.05) | 0.80 (0.05) | **0.73 (0.05)** | **0.68 (0.06)** |
|  | MG-GBLUP | **0.82 (0.05)** | **0.81 (0.05)** | **0.73 (0.05)** | **0.68 (0.06)** |
| Fruit weight | WPP | 0.52 (0.09) | 0.52 (0.09) | 0.57 (0.08) | 0.58 (0.08) |
|  | APP | 0.29 (0.11) | 0.3 (0.11) | 0.19 (0.10) | 0.2 (0.10) |
|  | Comb | **0.54 (0.10)** | **0.55 (0.09)** | **0.59 (0.08)** | **0.61 (0.07)** |
|  | MG-GBLUP | **0.54 (0.09)** | 0.54 (0.09) | **0.59 (0.08)** | 0.6 (0.08) |
| Fruit number | WPP | 0.48 (0.08) | 0.46 (0.08) | 0.5 (0.11) | 0.51 (0.11) |
|  | APP | 0.27 (0.12) | 0.24 (0.03) | 0.27 (0.11) | 0.29(0.11) |
|  | Comb | **0.49 (0.08)** | **0.47 (0.09)** | **0.52 (0.12)** | **0.54 (0.11)** |
|  | MG-GBLUP | 0.47 (0.08) | 0.45 (0.09) | 0.51 (0.11) | 0.52 (0.11) |

For the within-populations predictions, the predictive abilities varied between 0.33 and 0.75 for the elite material of the FBo-Hi dataset (respectively between 0.46 and 0.79 for the REFPOP dataset), and between 0.34 and 0.8 for the genetic resources of the FBo-Hi dataset (respectively between 0.5 and 0.79 for the REFPOP dataset). In both datasets, predictive abilities were high for harvest date (between 0.69 and 0.8 for FBo-Hi and between 0.6 and 0.79 for REFPOP, see Figure 3A) and fruit over-color (between 0.61 and 0.75 for FBo-Hi and between 0.64 and 0.79 for REFPOP, see Figure 3B), while they were moderately high for the other traits (0.33 to 0.58). The predictive abilities were higher when predicting harvest date (with a difference up to 0.19 and smaller standard errors), fruit weight (difference up to 0.06) and fruit number (difference up to 0.05) in the genetic resources than in the elite material, while fruit over-color (difference up to 0.15) and juiciness (difference up to 0.09) were better predicted when using elite material in the training and validation set.

**Figure 3.**
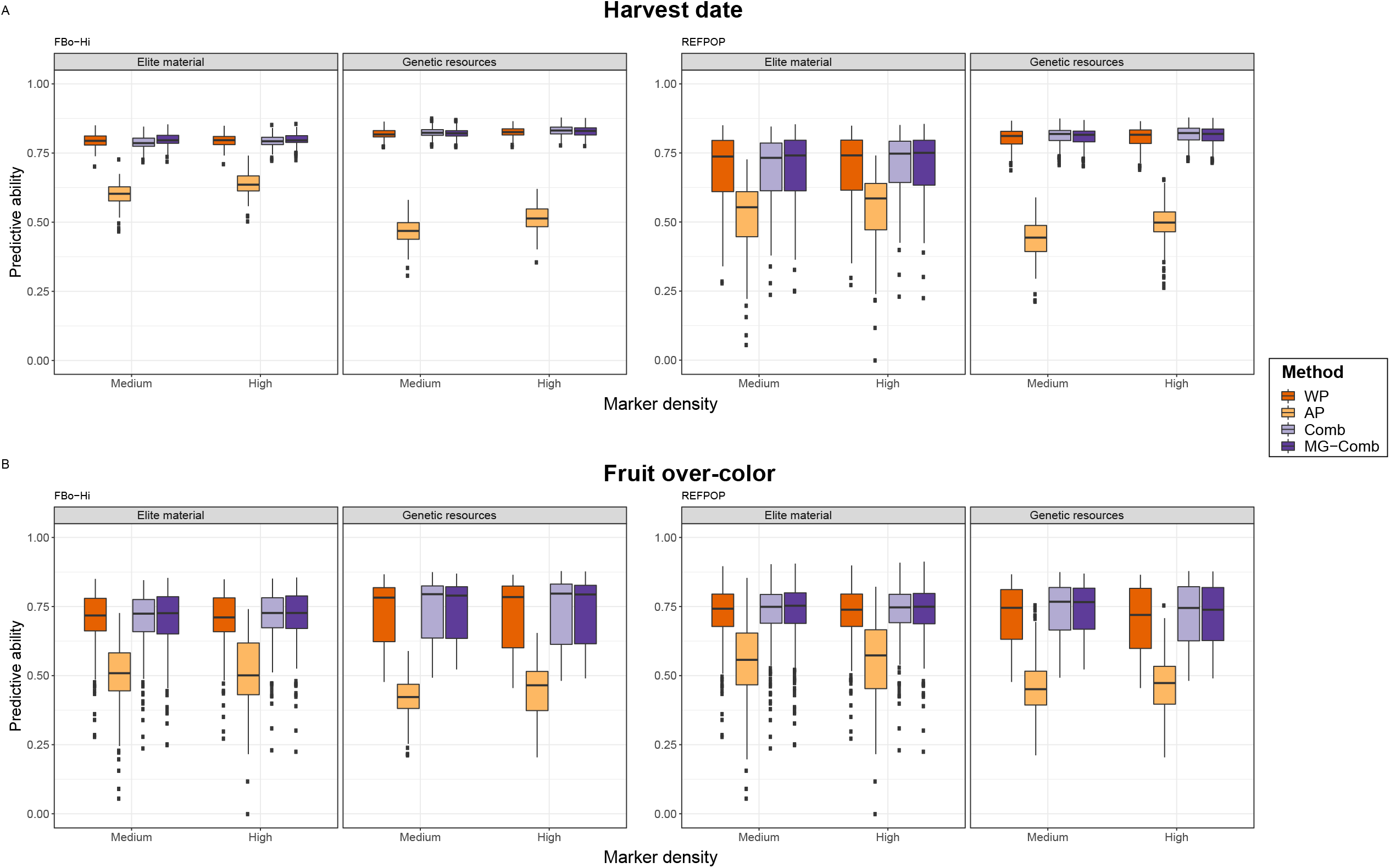
Predictive abilities for harvest date and fruit over-color in the FBo-Hi or REFPOP dataset with medium and high marker density. **WP**: within-population prediction **AP**: across-population prediction **Comb**: combination prediction **MG-Comb**: combination prediction with the MG-GBLUP method

The across-population predictions always resulted in lower predictive abilities than the within-population predictions, with values ranging from 0.13 (juiciness) to 0.60 (fruit over-color) for the FBo-Hi dataset and from 0.19 (fruit weight) to 0.79 (fruit over-color) for the REFPOP dataset. The standard errors of predictive abilities were also generally higher than for within-population predictions. Across-population predictive abilities could be moderately high for some traits, such as harvest date (between 0.44 and 0.54 for the FBo-Hi dataset and between 0.41 and 0.47 for the REFPOP dataset) or fruit over-color (between 0.43 and 0.60 for the FBo-Hi dataset and between 0.52 and 0.73 for the REFPOP dataset). All the traits were better predicted when using genetic resources in the training set to predict elite material than when using elite material to predict genetic resources, with the exception of fruit number and crispness that were slightly better predicted in the latter case. The decrease in predictive ability between within and across-population prediction was also more important when genetic resources constituted the validation set, especially for harvest date (for instance, with a decrease from 0.79 to 0.41 for the REFPOP dataset at medium marker density), fruit weight (from 0.57 to 0.19) and juiciness (from 0.49 to 0.18).

For all the studied traits in the two datasets, combining populations into a unique training set allowed to obtain predictive abilities that were slightly higher than with the corresponding within-population prediction, except when predicting the harvest date of the elite material with a medium density array in both datasets, which led to the same predictive ability. The highest increases in predictive ability were observed for the fruit over-color in the genetic resources in both datasets (from 0.64 to 0.70 in the FBo-Hi dataset and from 0.69 to 0.73 in the REFPOP dataset with medium density). Allowing marker effects to differ between populations (MG-GBLUP model) never led to improvements of the predictive ability compared to a prediction using a standard GBLUP model, except for juiciness when predicting genetic resources (increase of 0.02 at both medium and high marker density) and fruit over-color when predicting elite material (increase of 0.01 at both medium and high marker density).

Using a high marker density allowed higher predictive abilities compared to medium density only for harvest date and fruit weight, especially when predicting across populations (for instance, an increase from 0.44 to 0.49 for the FBo-Hi dataset and from 0.41 to 0.47 for the REFPOP dataset was observed for harvest date when predicting genetic resources with a training set of elite material). For some traits, measured predictive abilities were higher with medium marker density than with high marker density, as can be observed for over-color in both datasets and for fruit acidity. The decrease in predictive ability was also more important for across-population predictions for these two traits, especially when predicting genetic resources (decrease of 0.05 for both traits when predicting with the Fbo-Hi dataset and of 0.04 for acidity with the REFPOP dataset). For the remaining traits, the influence of the marker density depended on the population used in the validation set: predictive abilities for fruit number, juiciness and crispness decreased when predicting elite material using a high marker density but increased in the genetic resources.

### Proportion of the combined populations

When predicting the hybrids, we also investigated the influence of the proportion of elite material and genetic resources used in the training set. Four methods were compared to define the training set: the CDpop algorithm (CDstrat method), a choice based on kinship between the training set and validation set (Kinship method), or a random choice of genotypes either within each population (Random_strat method) or regardless of the populations (Random method). The results for harvest date and fruit over-color are presented in Figures 4-5 and the predictive abilities for fruit weight and fruit number are presented in Figures S14-15.

**Figure 4.**
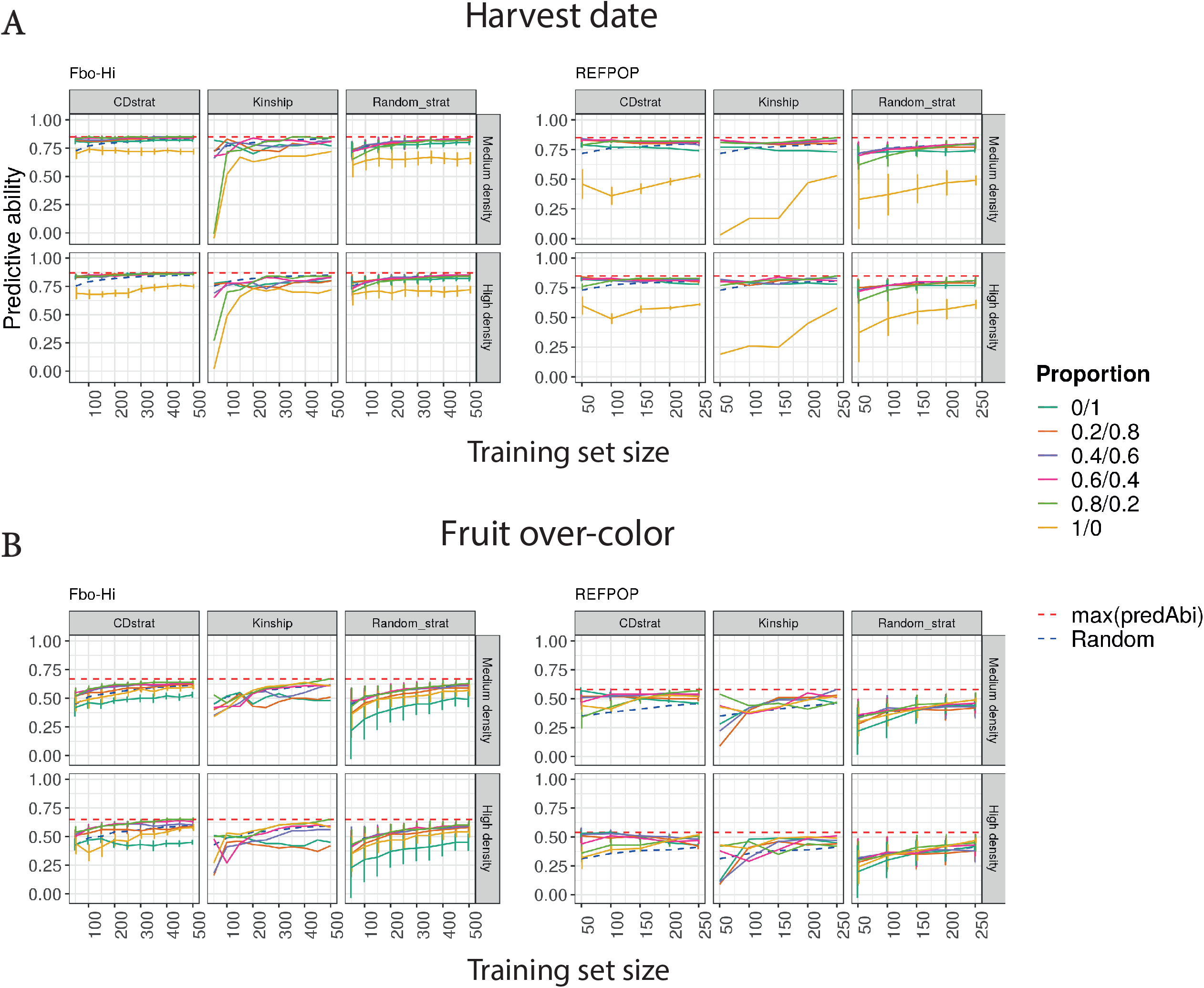
Predictive abilities for harvest date and fruit over-color in the dataset of hybrids with medium and high marker density when the training set is composed of varying proportions of elite material and genetic resources of the FbBo-Hi or REFPOP dataset. **max(predAbi)**: maximum predictive ability obtained for a given marker density with one of the three tested methods regardless of the training set size **Random**: predictive ability obtained when randomly choosing the genotypes included in the training set

We observed that the predictions based on the CDstrat method generally outperformed the Kinship, Random_strat and Random methods (with the exception of the prediction of the fruit number where the Random_strat method allowed to reach slightly higher predictive abilities). The standard errors of the predictive abilities were also lower when using the CDstrat method than when using the Random_strat or Random method. This observation holds true for all the tested training population sizes and at medium and high marker density in the case of harvest date predicted with the FBo-Hi dataset and for fruit weight (except for one tested proportion, see below). When predicting harvest date with the REFPOP dataset, the CDstrat and Kinship methods gave very similar results for all the tested proportions and training set sizes, but CDstrat allowed to reach higher predictive abilities for the 1/0 proportion, especially for small training set sizes (with a maximum predictive ability difference of 0.46 between the two methods for a TS of 50 genotypes). For fruit over-color, the best method depended on the tested proportion: with the FBo-Hi dataset, CDstrat outperformed Kinship except for the extreme proportions (i.e. TS constituted of only one of the two populations), the highest increases in predictive ability being again measured for small training set sizes. With the REFPOP dataset, CDstrat outperformed Kinship only until a TS size of 100 genotypes, and the two methods gave similar results for larger TS sizes. For all the tested TS sizes, most genotypes included in the TS with the CDstrat method were selected at least 15 times out of the 20 repetitions (data not shown).

The predictive abilities based on the CDstrat method rapidly reached a plateau for harvest date. The maximum predictive ability of 0.85 could be reached with 100 genotypes from the FBo-Hi dataset at medium density, and the maximum predictive ability was reached with 50 genotypes from the REFPOP dataset for both densities. For fruit over-color, a bigger TS size resulted in higher predictive abilities with the FBo-Hi dataset for all but the 0/1 and 1/0 proportions. The increase in TS size also led to an increase in predictive ability only for some proportions for fruit over-color with the REFPOP dataset: increasing the TS from 50 to 250 genotypes allowed an increase in predictive ability for proportions 0.4/0.6 to 1/0 (with increase between 0.02 for the 0.4/0.6 proportion and 0.43 for the 0.8/0.2 proportion) but the predictive ability decreased for the 0/1 and 0.2/0.8 proportions. A similar pattern was observed for fruit weight: when increasing the TS size, predictive abilities also increased for proportions 0.6/0.4 to 1/0 but decreased for proportion 0.2/0.8 and 0/1 beyond respectively 150 and 100 genotypes in the TS. The predictive abilities for fruit number increased when the TS size increased for all the tested proportions, but the gains were smaller than for the other traits (maximum increase from 0.16 to 0.26 when increasing the TS size from 50 to 250 genotypes). In all the studied traits, the standard errors of the predictive abilities also decreased when the TS size increased.

With the Kinship method, the largest increases in predictive ability were measured for harvest date predicted with the FBo-Hi dataset, especially for the 0.8/0.2 and 1/0 proportions at medium marker density (respectively from -0.01 and -0.05 with a TS size of 50 genotypes to 0.84 and 0.72 with a TS size of 500 genotypes) but not in the REFPOP dataset: the predictions with 50 genotypes in the TS led to better predictive abilities than a TS with 250 genotypes, except for the 0.8/0.2 and 0/1 proportions. The increase in TS size also led to higher predictive abilities for all the tested proportions for fruit number, with larger gains than for the CDstrat method (with a maximum increase from 0.13 to 0.29 when increasing the TS size from 50 to 250 genotypes). The predictive ability also increased for fruit weight and fruit over-color for all the tested proportions, except for the 0/1 proportion for fruit weight and the 0.8/0.2 proportion for fruit over-color when predicted with the REFPOP dataset, with maximum predictive ability observed with a TS size of 50 genotypes in both cases.

For each trait, the proportion allowing the highest predictive ability depended on the TS size and the method used to select the genotypes of the TS. For the three methods, the worst predictive abilities were obtained when only one population was present in the TS, with predictive abilities lower than the ones obtained from random sampling in the two populations: the 0/1 proportion led to the lowest predictive abilities for fruit weight, fruit number and fruit over-color when predicted with the FBo-Hi dataset while for harvest date, the predictions with the 1/0 proportion gave the lowest predictive abilities. Note that for fruit weight and fruit over-color, the 1/0 proportion also led to predictive abilities lower than the predictive abilities obtained with the Random method.

As was observed in the within and across-population predictions, the effect of the marker density on predictive ability depended on the trait: harvest date generally benefited from the higher marker density in the two datasets, especially when the TS size increased. Interestingly, predictive abilities at high marker density with the 1/0 proportion were lower than with medium marker density for small TS sizes when predicting with the FBo-Hi dataset, while the increase in predictive ability due to higher marker density was the highest for the 1/0 proportion when predicting with the REFPOP dataset, with increases ranging from 0.08 to 0.15 with the CDstrat method. Predictive abilities for fruit weight were higher at high density only for TS sizes up to 100 genotypes and medium density led to higher predictive abilities for larger TS sizes with the CDstrat method, except for the 0/1 proportion, in which case high density always gave better results (with increases in predictive ability ranging from 0.04 and 0.13). Predictive abilities measured from high marker density data for fruit over-color and fruit number were slightly lower than when using medium marker density for every method and every tested proportion, except for the 1/0 proportion for fruit over-color predicted with the REFPOP dataset with the Kinship method which allowed slightly higher predictive abilities until a TS size of 200 genotypes.

## DISCUSSION

One possible limitation to the implementation of genomic selection in fruit tree crops is the establishment of a training set large enough to allow accurate prediction of selection candidates because of the possible costs of maintaining and phenotyping trees. In this study, we first evaluated the potential of combining two different apple populations to increase the size of the training set by estimating the predictive ability in this case. We then investigated the effect of using subsets of the two populations to predict hybrids resulting from crosses between genetic resource and elite material. In each case, we compared the predictive abilities obtained using either medium or high SNP marker density.

### Predictive abilities within and across populations

When predicting within-population, the measured predictive abilities were moderate to high, ranging from 0.33 for fruit crispness to 0.8 for harvest date. These values were overall in line with previous studies in apple that predicted the same traits. For example, harvest date was well predicted in other studies performed on apple (Migicovsky *et al*. 2016; McClure *et al*. 2018; Jung *et al*. 2020; Minamikawa *et al*. 2021) with predictive ability values between 0.6 and 0.8, while prediction accuracies for fruit weight and fruit quality traits were generally low to moderate (Kumar *et al*. 2015; Migicovsky *et al*. 2016; McClure *et al*. 2018; Minamikawa *et al*. 2021), with predictive abilities not exceeding 0.5. An exception was the study from Kumar *et al*. (2012b) that reported predictive ability values over 0.7 for traits related to fruit quality, in a case where genotypes from seven full-sibs families derived from a 4 × 2 factorial design were randomly used for cross-validation, leading to a higher relatedness between training and population set than in our study.

Across-population predictions always resulted in predictive abilities lower than within-population predictions. This is an expected result as linkage disequilibrium, allele effects and allele frequencies are generally different between populations. In our case, the LD decay was indeed more rapid in the genetic resources than in the elite material. The allele frequencies were similar along the 17 chromosomes but differed for some genomic regions, which could be due to selection, thus leading to different allele effects estimates as discussed in further. For some traits like fruit crispness or juiciness, the different phenotypic distribution in the two populations could also explain the poor predictive abilities in across-population prediction. Roth *et al*. (2020) reported a similar result when predicting apple fruit texture of full-sibs families using a germplasm collection. The high genomic correlations between the two populations could have suggested that across-population predictions could perform as well as within-population prediction, which was not the case in our study. Similar results were observed by Lyra *et al*. (2018) and Rio *et al*. (2019), who hypothesized that this apparently contradictory observation could be explained by population-specific allele frequencies at the causal QTL. Such differences are expected in our case because of selection (see below). The moderate to high predictive abilities observed for some traits like harvest date and fruit over-color and the high genomic correlations between the two populations nevertheless suggest that marker effects are conserved to a certain extent between genetic resources and elite material. In the case of the FBo-Hi dataset, the lower predictive abilities observed in the across-population predictions could also be due to GxE interactions since the trees of the two populations were phenotyped by different assessors in different sites and years. Combining genotypes that were phenotyped in environments that differ from the environments of the validation set is not always detrimental for genomic predictions (Jarquin *et al*. 2016) but when updating the training set, such genotypes should be discarded and replaced by genotypes from the same environment when it becomes possible to do so.

### Potential of combining populations

Combining datasets can help increase predictive abilities in genomic prediction by allowing larger training populations, which is a key parameter for accurate predictions. However, the marker effects may be different between the combined populations, in which case the addition of new genotypes to the training set is not expected to improve predictive abilities (Lund *et al*. 2016). For instance, several studies in livestock found no increase in predictive ability when combining breeds (Hayes *et al*. 2009; Erbe *et al*. 2012) and adding genetically distinct individuals to the training population can even result in lower predictive abilities (Lorenz and Smith 2015). Multi-population genomic prediction models combining populations that have diverged for a few generations are then expected to lead to higher predictive abilities (de Roos *et al*. 2009; Wientjes *et al*. 2016) and the choice of the populations to combine should be based on knowledge about the genetic distance between the populations.

Several studies showed evidence of selection during apple domestication, e.g. for fruit size (Yao *et al*. 2015; Duan *et al*. 2017), fruit quality traits (Wedger *et al*. 2021), metabolite content (Khan *et al*. 2014) or disease-related genes (Singh *et al*. 2019). However, little information is available about the genomic consequences of the more recent apple improvement and the resulting genetic differences between genetic resources and modern cultivars. Figure S4 shows that the genotypes from the elite material are less acid, juicier and crispier than genetic resources, which can be a consequence of breeders’ choices since these fruit quality traits have been targeted for decades in apple breeding programs (Janick and Moore 1975; Liao *et al*. 2021). On average, elite material is also harvested later than genetic resources, most probably because fruits with a later ripening are more firm (Migicovsky *et al*. 2016) and can be stored longer (Nybom *et al*. 2008). Modern varieties also have a lower phenolic content than genetic resources in general (Ceci *et al*. 2021; Watts *et al*. 2021). A recent pedigree reconstruction study showed that the two populations are separated by approximately 5-10 generations (Muranty *et al*. 2020), which could result in the persistence of marker phase and effects across populations.

In this study, combining elite material and genetic resources always led to higher predictive abilities than within-population predictions, even though the gains were generally modest. In particular, the combination allowed higher predictive abilities in the FBo-Hi dataset, in spite of the genotypes from the two populations being evaluated at different sites and years. This suggests that the potential adverse effect of GxE interactions on predictive ability were counterbalanced by the large increase in population size. The addition of genotypes from the alternative population did not decrease the predictive abilities when combining data, but we can hypothesize that selecting subsets of the two populations to combine closer genotypes into a training set could be advantageous. The approaches proposed in the Prop_hybrids scenario could thus be used for the Comb scenario. To further increase the size of the training population, historical data from trees that only have pedigree information could also be used in a single-step GBLUP model that uses both genotyped and non-genotyped individuals at the same time (Sood *et al*. 2020).

### Towards improvements of the predictive abilities

For all the studied traits, the MG-GBLUP model, which allows different but correlated effects between populations, did not perform better than the standard GBLUP model. This result can be explained in the light of the genomic correlations between the two populations: when the genomic correlation between the combined populations is equal to one, the MG-GBLUP is equivalent to the standard GBLUP. Therefore, the MG-GBLUP and standard GBLUP were expected to yield similar results since the genomic correlation between the genetic resources and elite material were high for all the traits. In addition, the MG-GBLUP shows limitations when dealing with genotypes that cannot clearly be assigned to a given population (Lehermeier *et al*. 2015), which is sometimes the case with the elite material and genetic resources (Figure 2 and Figure S8). In this case, genomic prediction models that account for admixture could be better adapted (Rio et al., 2020).

Ibañez-Escriche et al. (2009) suggested that models that fit population-specific marker effects may not be necessary at high marker density because a density high enough could allow the marker-QTL association to be the same in the combined populations. In this study, we observed limited increases in predictive abilities when using high marker density marker data. Other studies also reported that predictive abilities could reach a plateau past a given number of markers (Hickey *et al*. 2014; Jung *et al*. 2020) and that high marker density was not always needed to obtain high predictive abilities. For some traits, the predictive abilities were even lower than with medium marker density data. Two of these traits, namely fruit over-color and acidity, are controlled by an oligogenic determinism with a few known major genes (Chagné et al., 2007, Verma et al., 2019). As the GBLUP and MG-GBLUP models make the hypothesis that all the marker effects are drawn from the same normal distribution and do not allow marker effects to be null, a high marker density could overshrink the marker effects and poorly capture the effect of the large QTLs, as suggested by Daetwyler et al. (2010). Similar to our case, Erbe *et al*.(2012) found that predictive abilities were lower with a GBLUP model and 800K markers than with the same model with 50K markers and suggested that the number of effects to estimate was too important compared to the number of phenotypic records. In their case, models that allow marker effects to be set to zero or models resulting in a selection of a subset of markers led to higher predictive abilities. Bayesian models could also result in higher predictive abilities for traits with an oligogenic genetic architecture (Hayes et al., 2009). If major QTLs are known, they could also be considered as fixed effects (Bernardo, 2013, Sarinelli et al. 2019) or be weighted accordingly in the genomic relationship matrix (Tiezzi and Maltecca 2015; Raymond *et al*. 2018). Since genomic predictions are generally used for traits with a polygenic determinism, high marker density should give better results than lower marker densities in general. Moreover, DoVale *et al*. (2021) showed that, in outbred crops, genomic prediction models should be updated more regularly when high marker densities were not used. Using high marker density for genomic selection could thus prove useful when combining populations, especially if a high-quality imputation step can be implemented to reduce the costs of genotyping (Song *et al*. 2019).

### Optimization of the training set composition

When predicting the hybrids, we never observed a particular proportion of the two combined populations that would allow higher predictive abilities. However, using only one of the two populations was detrimental for each studied trait, regardless of the training set size or the method used to choose the genotypes. This point highlights the need to use at least some genotypes from both populations in the training set in order to achieve high enough predictive abilities. This observation can have at least two explanations. First, we observed in our data that the phenotypic variation in the hybrids is larger than the variation of the two populations taken separately. When using only one of the two populations in the training set, the range of phenotypic values of the chosen population can by consequence not be large enough to accurately estimate the marker effects. Second, if some alleles linked to the desired trait are in low frequency in one population and segregate in the other population, using only the population with the low frequency alleles as the training set will lead to incorrect marker effect estimations. For example, Migicovsky *et al*. (2021) showed that the *NAC18*.*1* gene marker, which is linked to harvest date and fruit firmness, is homozygous for the favorable allele in the nine most marketed varieties in the United-States, and several marker alleles detected by GWAS in the REFPOP panel are fixed in the elite population whereas they segregate in the genetic resources (Jung *et al*., submitted). As expected in this case, the prediction for harvest date in the hybrids when using only the elite material in the training set led to predictive abilities lower than when using both populations. Another such example would be the *Rvi6* gene that confers resistance to apple scab and all QTLs in the associated introgressed segment: the favorable allele at *Rvi6* segregates in elite material (Laurens 1998) but is absent in old varieties, because the gene introgression from the wild relative *M. floribunda* is recent (Gessler and Pertot 2012).

When the training set size is a limitation for the implementation of genomic selection, methods to optimize the choice of the genotypes in the TS when genomic data are already available can be advantageous. In this study, we observed that for three out of four traits the hybrids were better predicted when the TS was chosen based on the CDpop criterion compared to a choice based on kinship or genotypes sampled randomly, especially for small training set sizes. The CD algorithm has been implemented in order to better sample the genetic diversity than algorithms based on kinship alone (Rincent *et al*. 2012), which could explain the better performance of the CDstrat method, since it is probably necessary to use a training set with a large diversity to predict the hybrids, as discussed above.

Note that we applied the CD algorithm to each population separately in order to simultaneously study the effect of the proportion of the two populations on predictive abilities. One way to further optimize the composition of the training set could be to use the CDpop criterion with genotypes of the elite material and genetic resources in a single step, letting the algorithm choose the proportion of the two populations to be used. However, using the optimization procedure based on the CDpop criterion is computationally demanding, as the computation of the criterion requires to calculate the inverse of the genomic relationship matrix of the genotypes in the training and validation sets (Rincent *et al*. 2017), which could not be achieved in our case. It would also be interesting to evaluate the effect of the optimization methods proposed in this study in a case where the hybrids would be predicted by a set of hybrids. This situation should lead to the highest predictive abilities, and while the combination of populations proposed in this study is a suitable approach to initiate the prediction of hybrids, a training set composed of hybrids only should be envisioned as soon as enough phenotypic and genotypic data for the hybrids is available. Such a training set can be built gradually, by replacing a part of the genotypes of the two populations by newly phenotyped and genotyped hybrids (Fritsche-Neto *et al*. 2021).

## Conclusion

We showed in this study that combining genetic resources and elite material into a unique training set could be beneficial for genomic predictions. First, larger training populations can be obtained with this approach, leading to higher predictive abilities in return. Second, using both populations in the training set appeared necessary to predict “genetic resources x elite” hybrids. Combining populations could thus be an effective way to initiate breeding programs that incorporate genomic prediction when a large training population is too costly. The training set composition can be further optimized to reduce the number of genotypes to include, which can be of particular interest for traits that are hard or long to phenotype, like biennial fruit bearing or abiotic stress tolerance. The proposed approach can be used in other fruit tree species provided that the genetic differences of the combined populations are taken into account when necessary.

## Data availability

All SNP genotypic data generated with the 480K array used in this study have been deposited in the INRAe dataset archive (https://data.inrae.fr/) at https://doi.org/10.15454/IOPGYF. The SNP genotypic data of the REFPOP elite material generated with the 20K array and used in this study have been deposited in the INRAe dataset archive at https://doi.org/10.15454/1ERHGX. The SNP genotypic data of the elite material of the FBo-Hi panel generated using the 20K array and used in this study will be deposited in the same INRAe dataset archive upon acceptance of the manuscript. The SNP genotypic data of the hybrids generated using the 20K array in this study will be deposited in the same INRAe dataset archive upon acceptance of the manuscript. The raw phenotypic data of the REFPOP panel will be deposited in the INRAe dataset archive upon acceptance of Jung et al, submitted, paper. The raw phenotypic data of the genetic resources of the FBo-Hi dataset, the BLUPs of clonal values of the elite material of the FBo-Hi dataset, the adjusted phenotypic data of the hybrids will be deposited in the INRAe dataset archive upon acceptance of the present manuscript. The scripts used to obtain the results presented in this manuscript are available at https://sourcesup.renater.fr/projects/apple-gensel.

## Acknowledgments

We would like to thank the Horticole Experimental Unit (UE Horti) for maintaining and phenotyping the trees that served in this study, as well as Sandra Gabard for her help with the phenotyping of the hybrids in Angers and the collaborators of the REFPOP network for evaluating trees and reporting phenotypic data of the REFPOP panel. The authors in particular thank Michaela Jung for collecting and cleaning the raw data from the different partners of the project.

