## Supplemental tables and figures for "Combining genetic resources and elite material populations to improve the accuracy of genomic prediction in apple"

**Table S1**. Mating design and family sizes of the hybrids panel

| | | Genetic resources | | | | |
|---|---|---|---|---|---|---|
| | | X08233 | X08483 | X02353 | X02640 | X08488 |
| Elite | X02437 | 65 | | | | |
| | X03263 | | | | 39 | |
| | X03318 | | | | | 49 |
| | X06407 | | 38 | | | |
| | X06683 | | | 50 | | |
| | X06963 | 31 | | | | |
| | X07860 | 51 | | | | |
| | X08151 | 44 | 50 | | | |
| | X08486 | 56 | | | | |

**Table S2**. Year range and heritability of the traits measured in the FBo-Hi dataset

| | **CRA-W** | | **INRA** | | **NFC** | | **RBIPH** | | **SLU** | | **UNIBO** | |
|---|---|---|---|---|---|---|---|---|---|---|---|---|
| | No. years (range) | $H^2$ | No. years (range) | $H^2$ | No. years (range) | $H^2$ | No. years (range) | $H^2$ | No. years (range) | $H^2$ | No. years (range) | $H^2$ |
| Acidity | 13 (1986-2014) | 0.90 | 8 (2002-2014) | 0.80 | 1 | - | 2 (2006-2010) | 0.90 | 3 (2012-2014) | 0.90 | 5 (1988-2014) | 0.7 |
| Crispness | 8 (1987-2013) | 0,80 | 4 (2010-2014) | 0.70 | 2 (2012-2013) | 0.59 | 3 (2012-2014) | 0.72 | 2 (2012-2014) | 0.97 | 3 (2012-2014) | 0.5 |
| Juiciness | 12 (1986-2013) | 0,76 | 7 (2004-2014) | 0.70 | 2 (2012-2013) | 0.58 | 5 (2006-2010) | 0.74 | 3 (2012-2014) | 0.89 | 3 (2012-2014) | 0.6 |
| Fruit over-color | 4 (1989-2013) | 0,90 | 7 (2002-2010) | 0.81 | 1 | - | 5 (2006-2010) | 0.93 | 3 (2012-2014) | 0.94 | 9 (1987-2014) | 0.94 |
| Harvest date | 1 (2013) | - | 3 (2012-2014) | 0.96 | 3 (2012-2014) | 0.88 | - | - | 3 (2012-2014) | 0.98 | 3 (2012-2014) | 0.97 |

**Table S3**. Pairwise $F_{ST}$ values computed from marker data for the elite material, genetic resources and hybrids in the two datasets

| Fbo-Hi dataset | | |
|---|---|---|
| Population | Elite material | Genetic resources |
| Genetic resources | 0.023 | - |
| Hybrids | 0.029 | 0.017 |
| **REFPOP dataset** | | |
| Population | Elite material | Genetic resources |
| Genetic resources | 0.021 | - |
| Hybrids | 0.029 | 0.017 |

**Figure S1.** Linkage disequilibrium decay for SNPs within a 500kb distance in the elite material and genetic resources of the Fbo-Hi and REFPOP datasets

**Figure S2.** Frequence of the minor allele in the genetic resources and of the same allele in the elite material along the 17 chromosomes of the apple genome in the FBo-Hi dataset. Allele frequencies are computed using sliding windows of 2Mb with a shift of 400kb.

**Figure S3.** Frequence of the minor allele in the genetic resources and of the same allele in the elite material along the 17 chromosomes of the apple genome in the REFPOP dataset. Allele frequencies are computed using sliding windows of 2Mb with a shift of 400kb.

**Figure S4.** Phenotypic distribution of the traits measured in the FBo-Hi dataset only. **A**. Acidity **B.** Juiciness **C.** Crispness

**Figure S5.** Phenotypic distribution of the traits measured in the REFPOP dataset and the dataset of hybrids. **A**. Fruit number **B.** Fruit weight

**Figure S6.** Phenotypic distribution of the traits measured in the dataset of hybrids and FBo-Hi and REFPOP datasets. **A**. Harvest date **B.** Fruit over-color

**Figure S7.** Broad-sense heritability for each site and year in the REFPOP dataset. **BEL** Belgium **CHE** Switzerland **ESP** Spain **FRA** France **ITA** Italy

**Figure S8**. Principal Component Analysis (PCA) performed using pruned marker data of the REFPOP and hybrids panel.

**Figure S9.** Predictive abilities for acidity in the FBo-Hi dataset with medium and high marker density. **WP**: within-population prediction **AP**: across-population prediction **Comb**: combination prediction **MG-Comb**: combination prediction with the MG-GBLUP method

**Figure S10.** Predictive abilities for crispness in the FBo-Hi dataset with medium and high marker density. **WP**: within-population prediction **AP**: across-population prediction **Comb**: combination prediction **MG-Comb**: combination prediction with the MG-GBLUP method

**Figure S11.** Predictive abilities for juiciness in the FBo-Hi dataset with medium and high marker density. **WP**: within-population prediction **AP**: across-population prediction **Comb**: combination prediction **MG-Comb**: combination prediction with the MG-GBLUP method

**Figure S12.** Predictive abilities for fruit number in the REFPOP dataset with medium and high marker density. **WP**: within-population prediction **AP**: across-population prediction **Comb**: combination prediction **MG-Comb**: combination prediction with the MG-GBLUP method

**Figure S13.** Predictive abilities for fruit weight in the REFPOP dataset with medium and high marker density. **WP**: within-population prediction **AP**: across-population prediction **Comb**: combination prediction **MG-Comb**: combination prediction with the MG-GBLUP method

**Figure S14.** Predictive abilities for fruit number in the dataset of hybrids with medium and high marker density when the training set is composed of varying proportions of elite material and genetic resources of the REFPOP dataset. **max(predAbi)**: maximum predictive ability obtained for a given marker density with one of the three tested methods regardless of the training set size **Random**: predictive ability obtained when randomly choosing the genotypes included in the training set

**Figure S15.** Predictive abilities for fruit weight in the dataset of hybrids with medium and high marker density when the training set is composed of varying proportions of elite material and genetic resources of the REFPOP dataset. **max(predAbi)**: maximum predictive ability obtained for a given marker density with one of the three tested methods regardless of the training set size **Random**: predictive ability obtained when randomly choosing the genotypes included in the training set

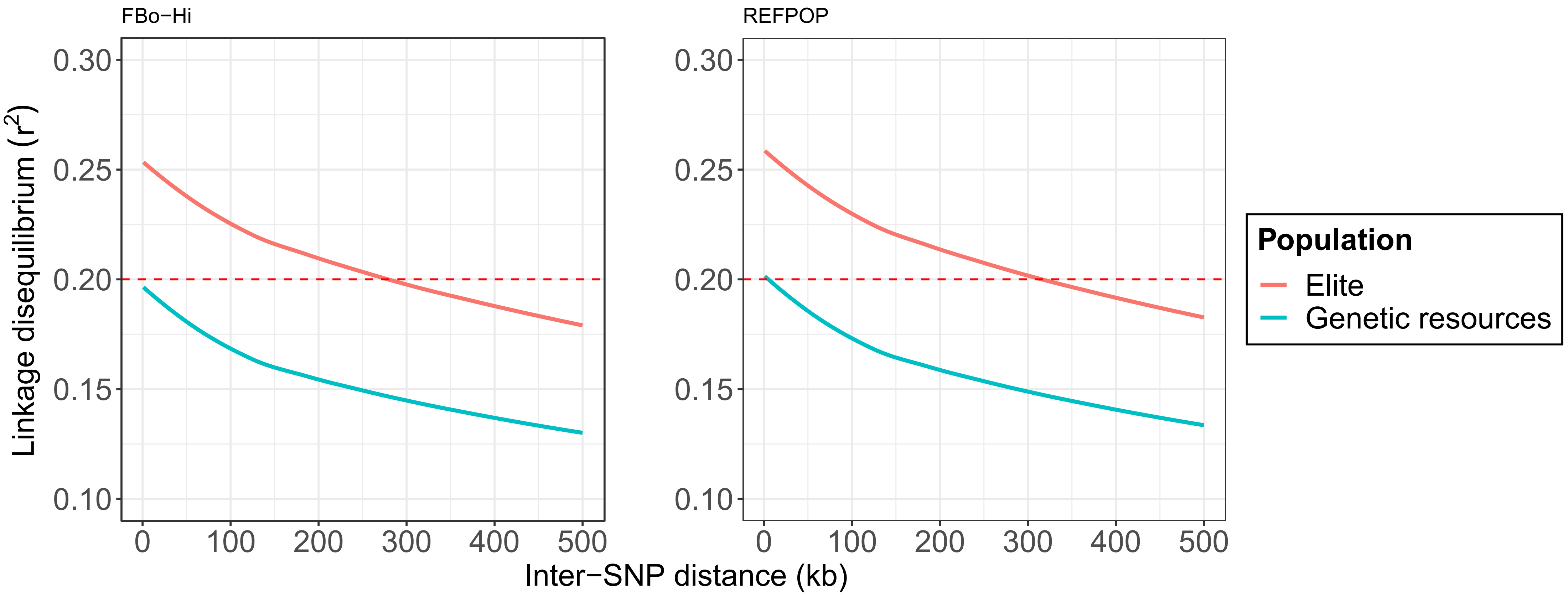

FBo−Hi
REFPOP
Linkage disequilibrium (r2)
0.30
0.25
0.20
0.15
0.10
0
100
200
300
400
500
Inter−SNP distance (kb)
Population
Elite
Genetic resources

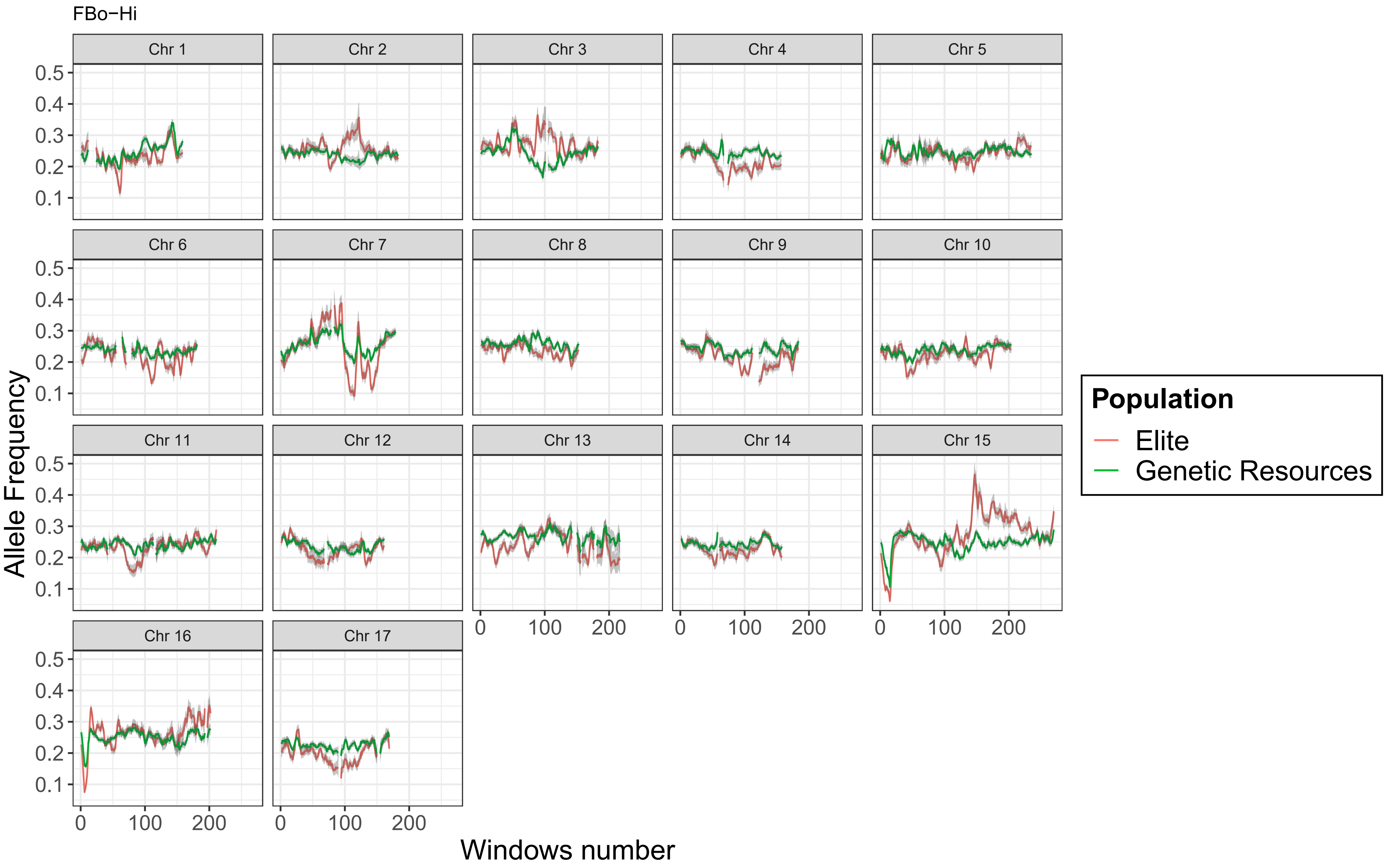
FBo−Hi
Chr 1
Chr 2
Chr 3
Chr 4
Chr 5
Chr 6
Chr 7
Chr 8
Chr 9
Chr 10
Chr 11
Chr 12
Chr 13
Chr 14
Chr 15
Chr 16
Chr 17
Allele Frequency
0.5
0.4
0.3
0.2
0.1
0
100
200
Windows number
Population
Elite
Genetic Resources

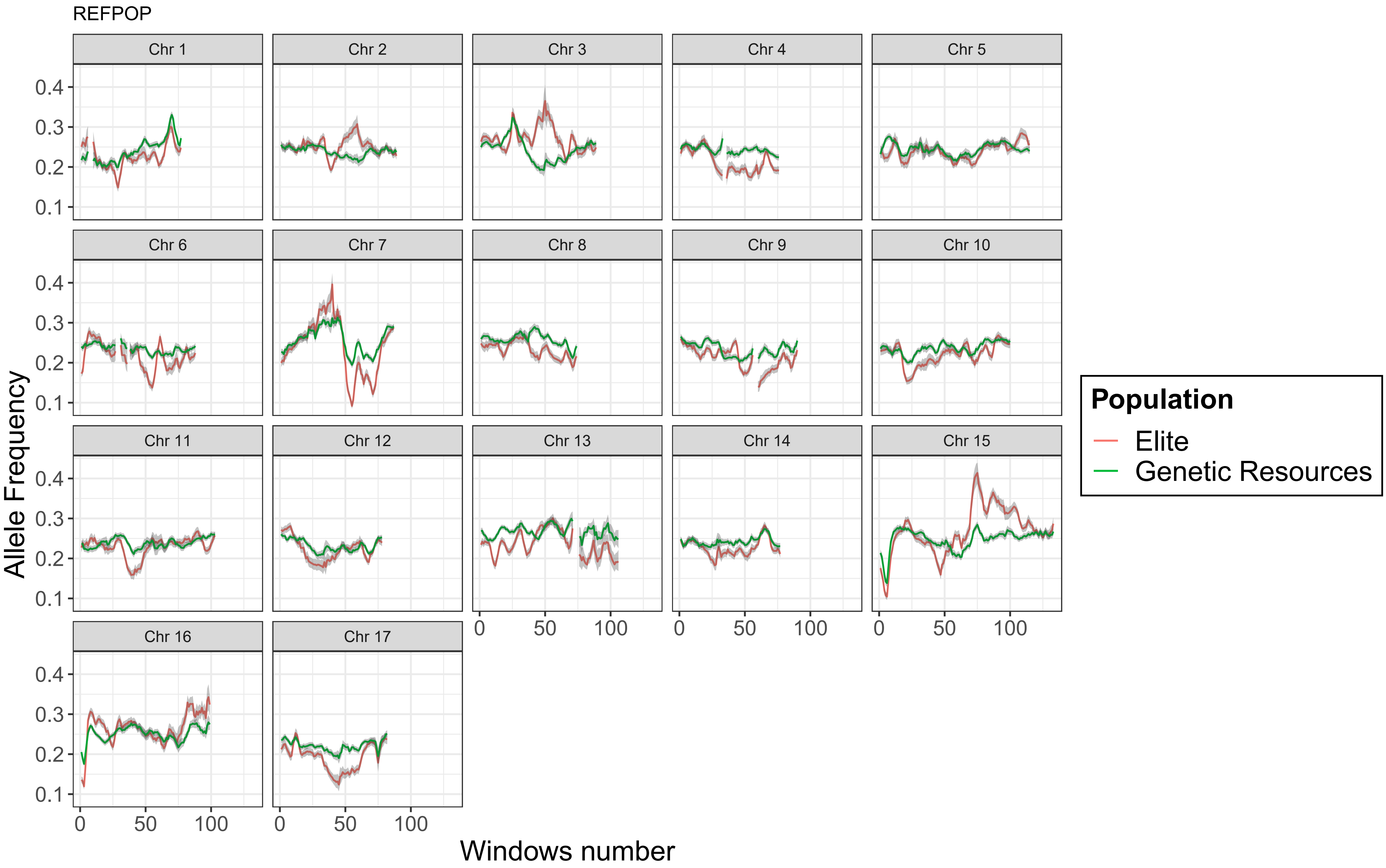
REFPOP
Chr 1
Chr 2
Chr 3
Chr 4
Chr 5
Chr 6
Chr 7
Chr 8
Chr 9
Chr 10
Chr 11
Chr 12
Chr 13
Chr 14
Chr 15
Chr 16
Chr 17
0.1
0.2
0.3
0.4
0
50
100
Allele Frequency
Windows number
Population
Elite
Genetic Resources

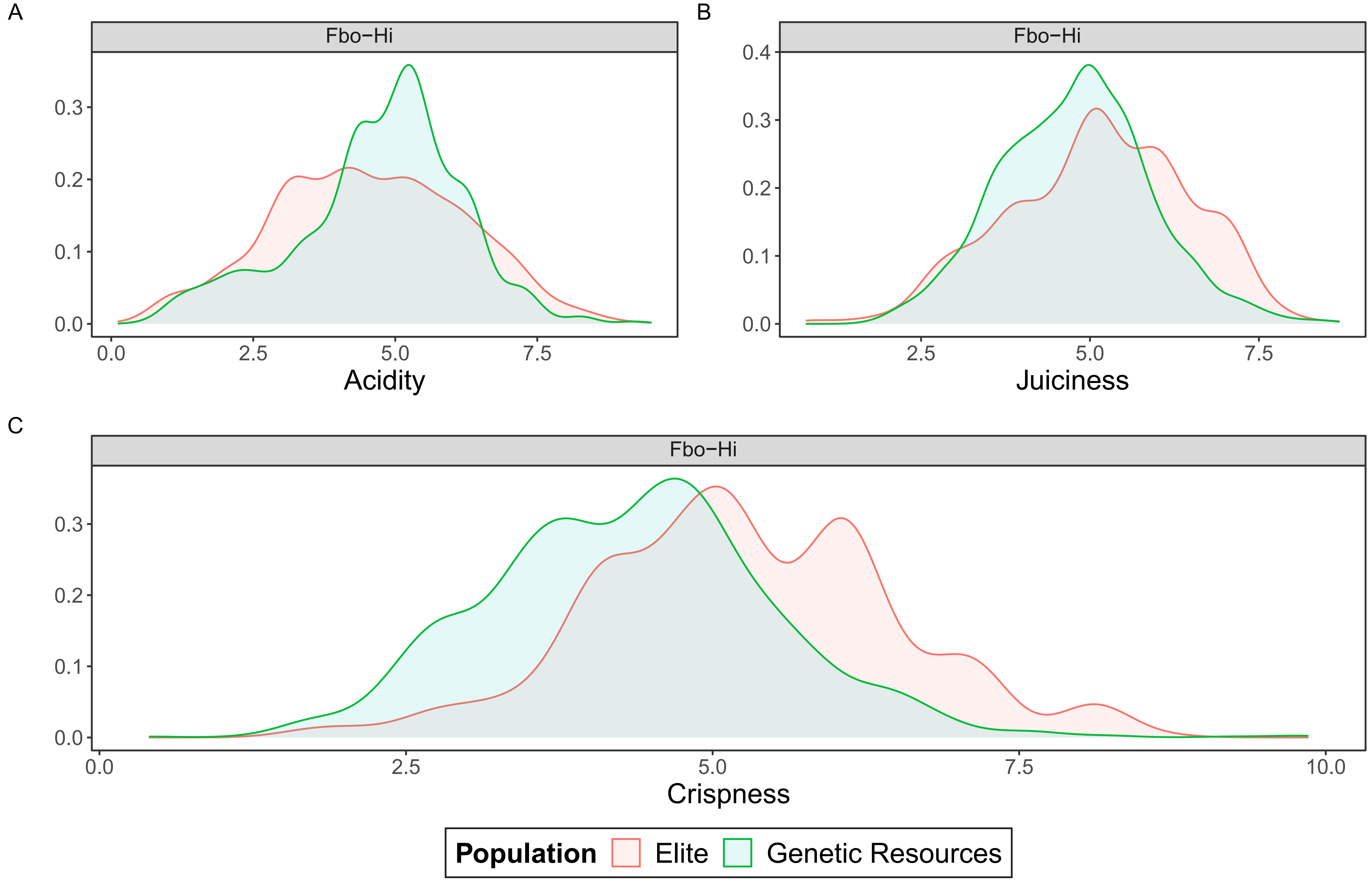

A
Fbo−Hi
0.3
0.2
0.1
0.0
0.0
2.5
5.0
7.5
Acidity
B
Fbo−Hi
0.4
0.3
0.2
0.1
0.0
2.5
5.0
7.5
Juiciness
C
Fbo−Hi
0.3
0.2
0.1
0.0
0.0
2.5
5.0
7.5
10.0
Crispness
Population
Elite
Genetic Resources

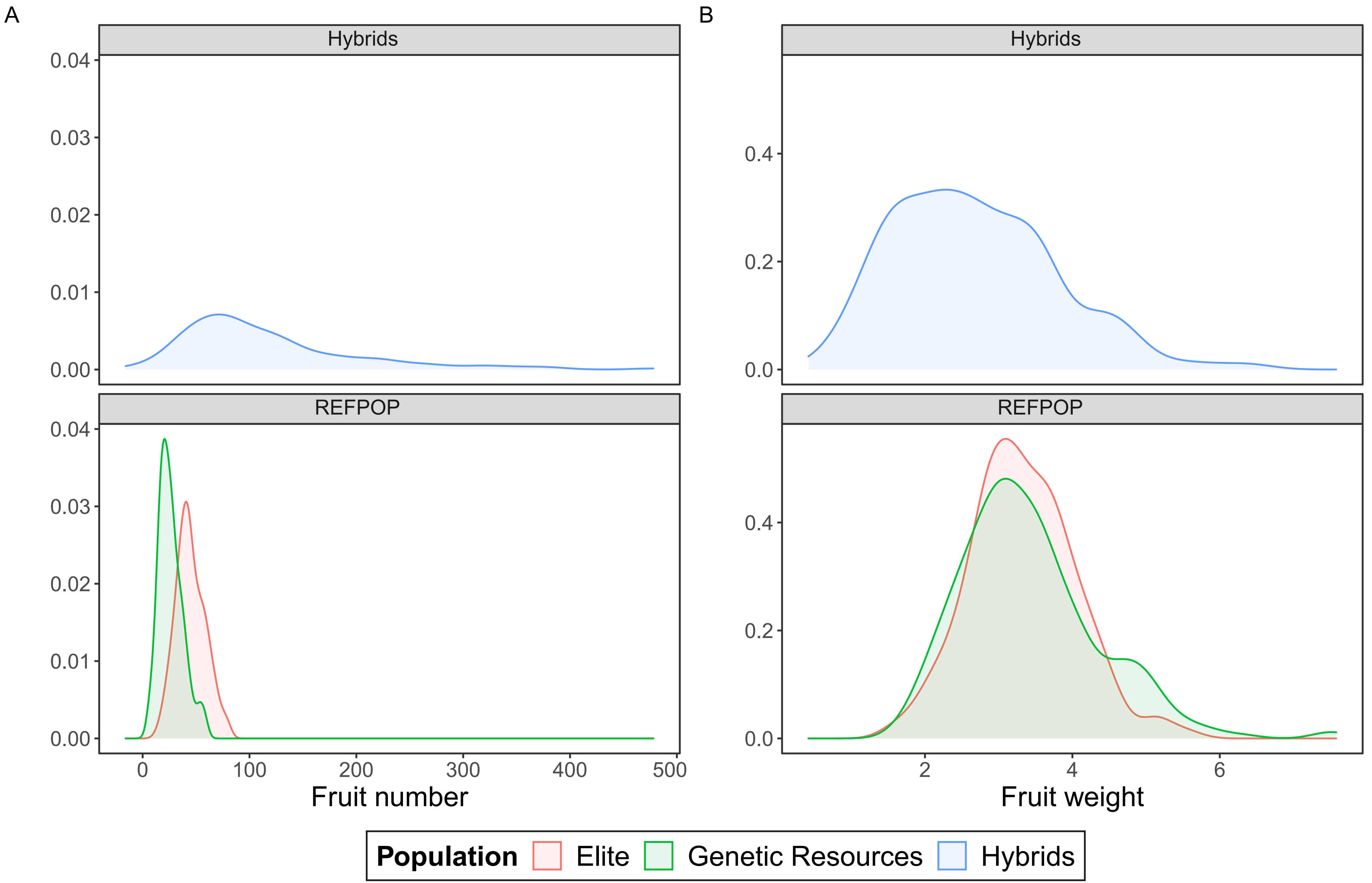

A
B
Hybrids
REFPOP
0.04
0.03
0.02
0.01
0.00
0
100
200
300
400
500
Fruit number
0.4
0.2
0.0
2
4
6
Fruit weight
Population
Elite
Genetic Resources
Hybrids

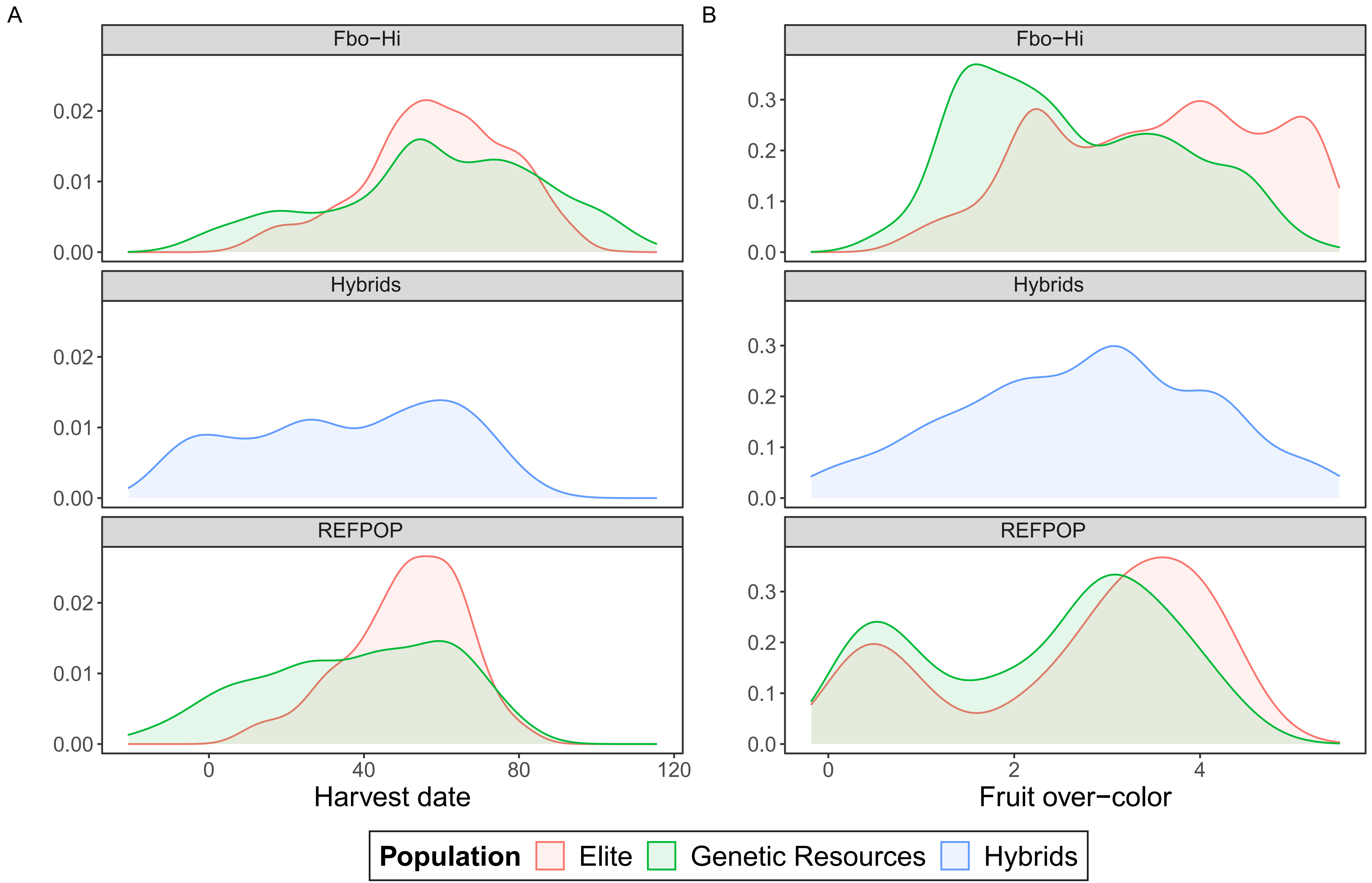
A
B
Fbo−Hi
Hybrids
REFPOP
0.00
0.01
0.02
0.0
0.1
0.2
0.3
0
40
80
120
2
4
Harvest date
Fruit over−color
Population
Elite
Genetic Resources
Hybrids

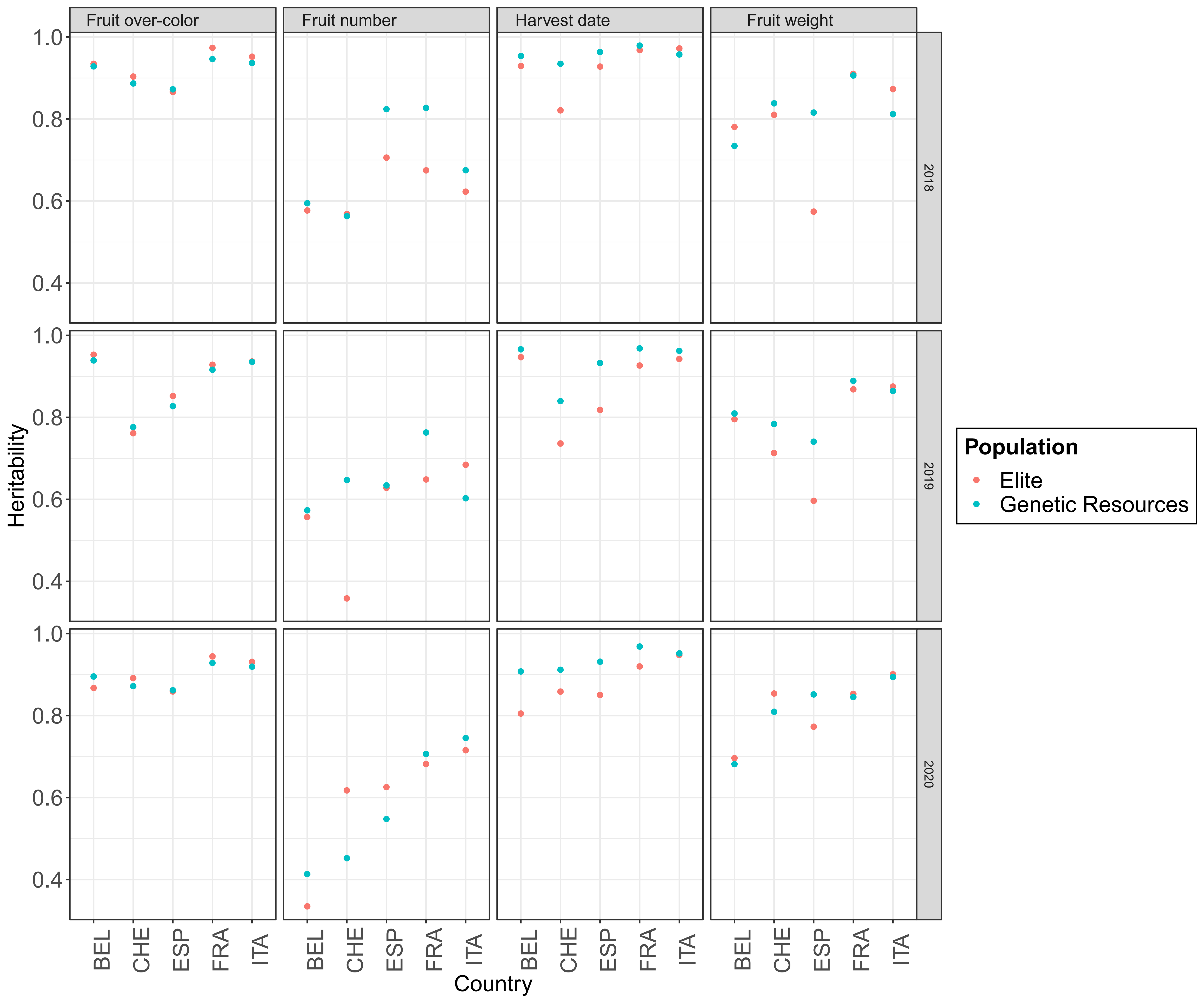

Fruit over-color
Fruit number
Harvest date
Fruit weight
2018
2019
2020
Heritability
1.0
0.8
0.6
0.4
BEL
CHE
ESP
FRA
ITA
Country
Population
Elite
Genetic Resources

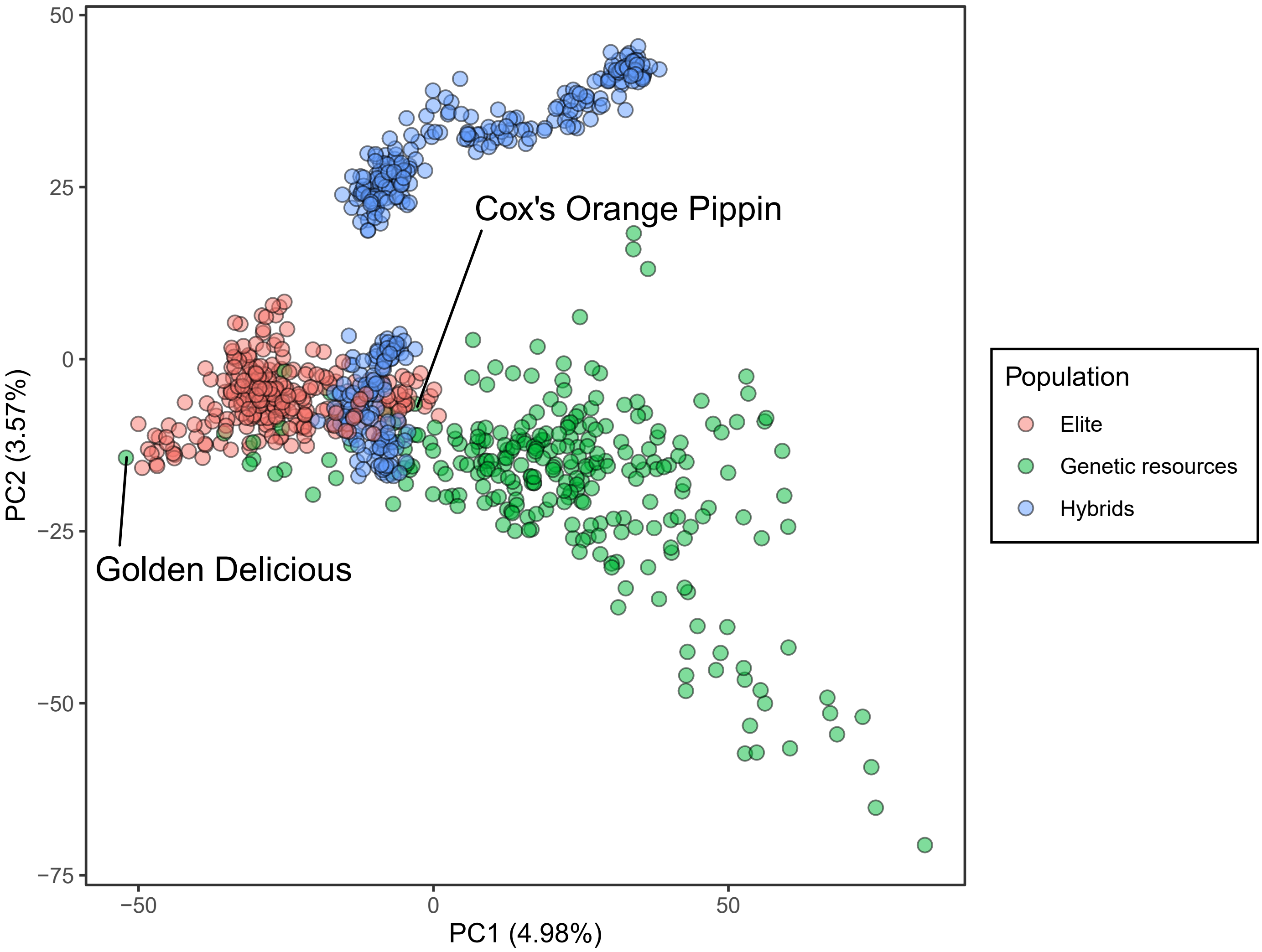

Cox's Orange Pippin
Golden Delicious
PC1 (4.98%)
PC2 (3.57%)
50
25
0
−25
−50
−75
−50
0
50
Population
Elite
Genetic resources
Hybrids

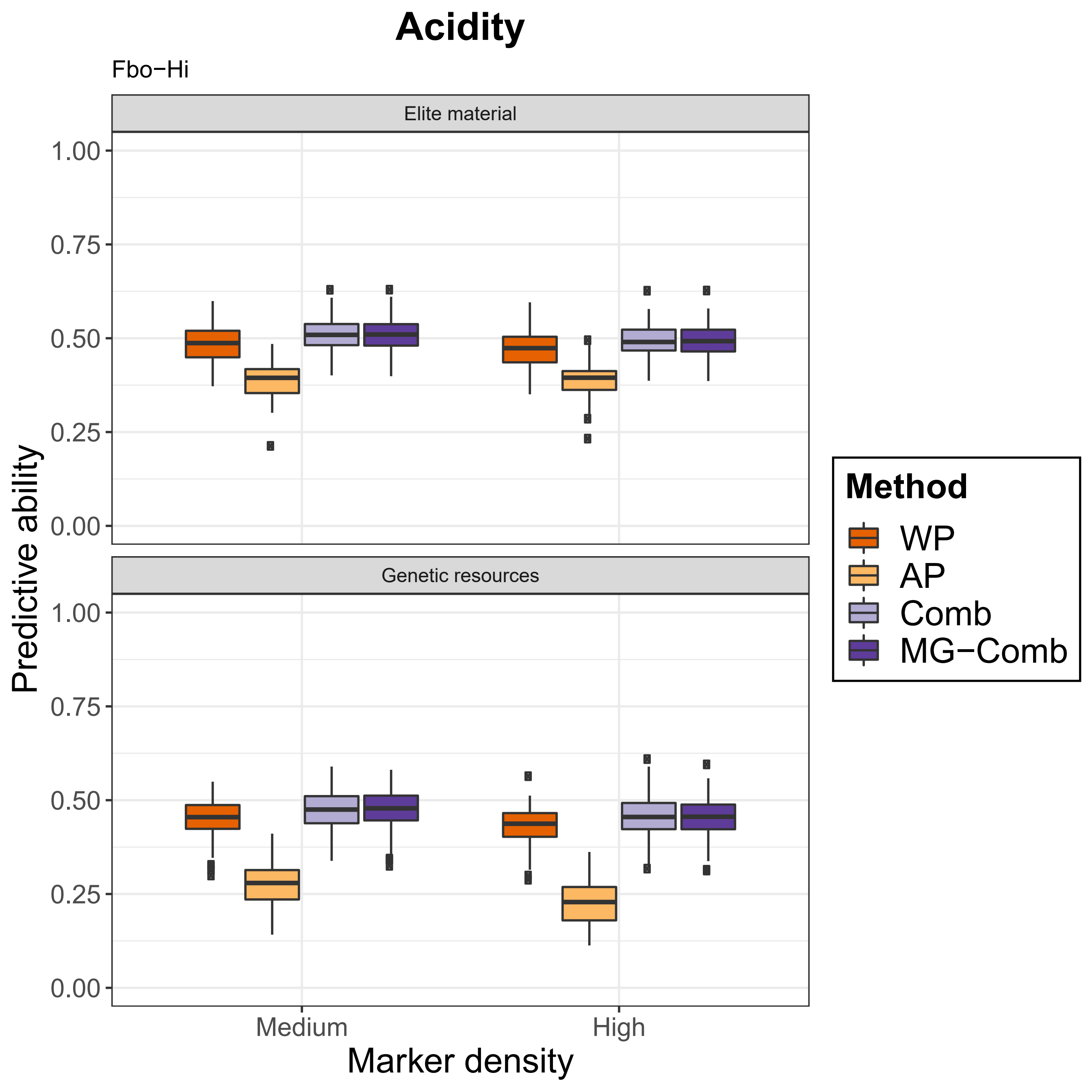
Acidity
Fbo−Hi
Elite material
Genetic resources
Predictive ability
0.00
0.25
0.50
0.75
1.00
Medium
High
Marker density
Method
WP
AP
Comb
MG−Comb

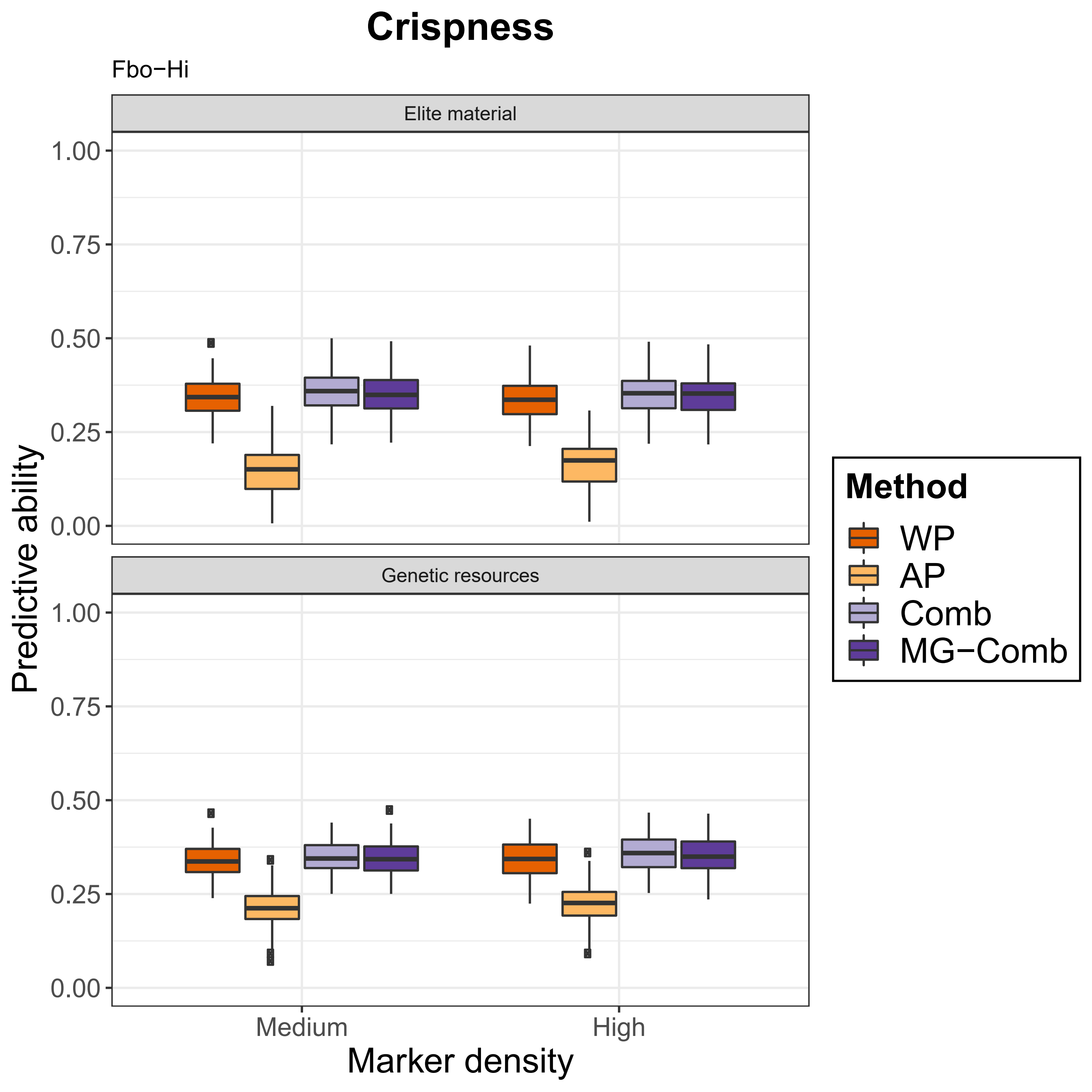

Crispness
Fbo−Hi
Elite material
Genetic resources
1.00
0.75
0.50
0.25
0.00
Predictive ability
Medium
High
Marker density
Method
WP
AP
Comb
MG−Comb

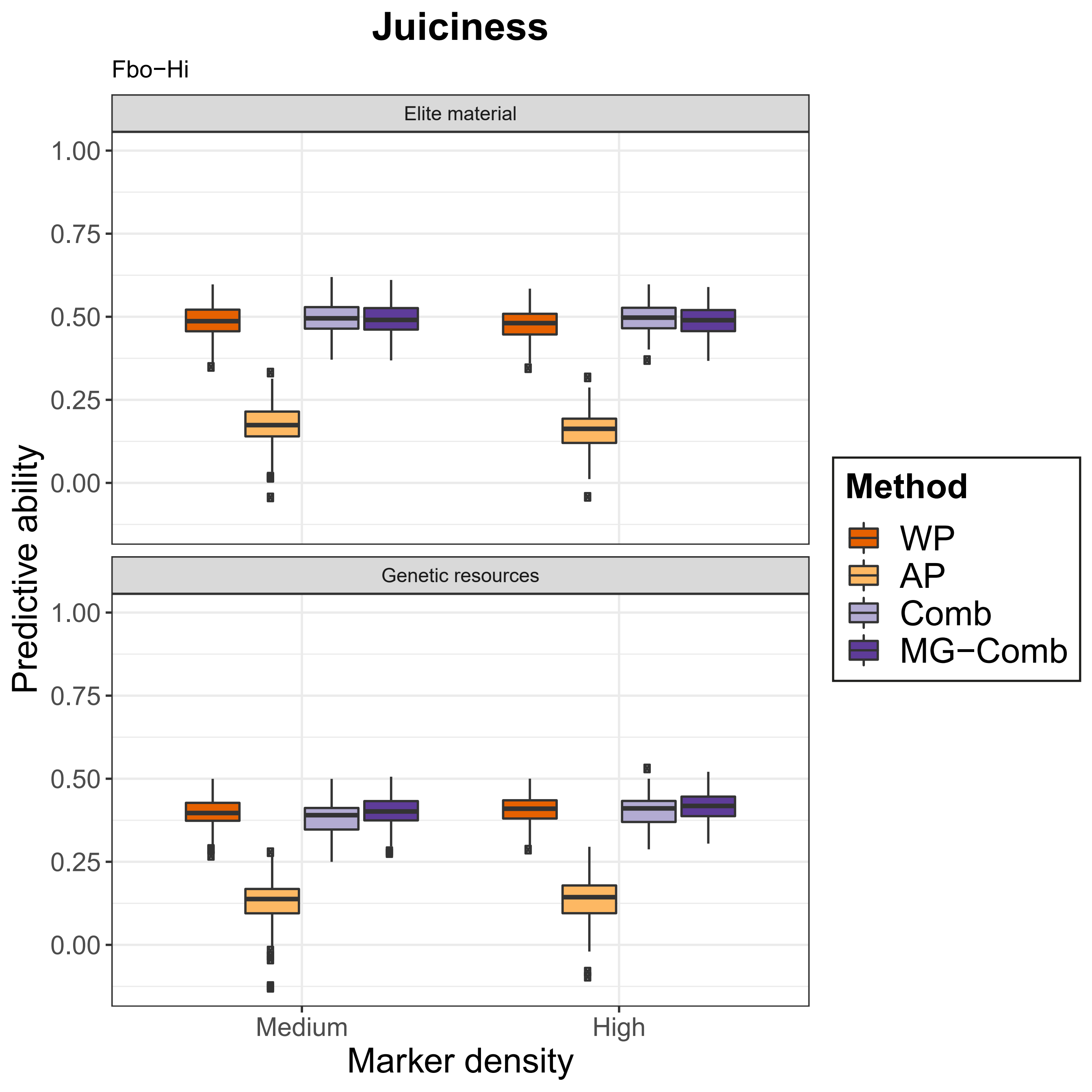

Juiciness
Fbo−Hi
Elite material
Genetic resources
1.00
0.75
0.50
0.25
0.00
Predictive ability
Medium
High
Marker density
Method
WP
AP
Comb
MG−Comb

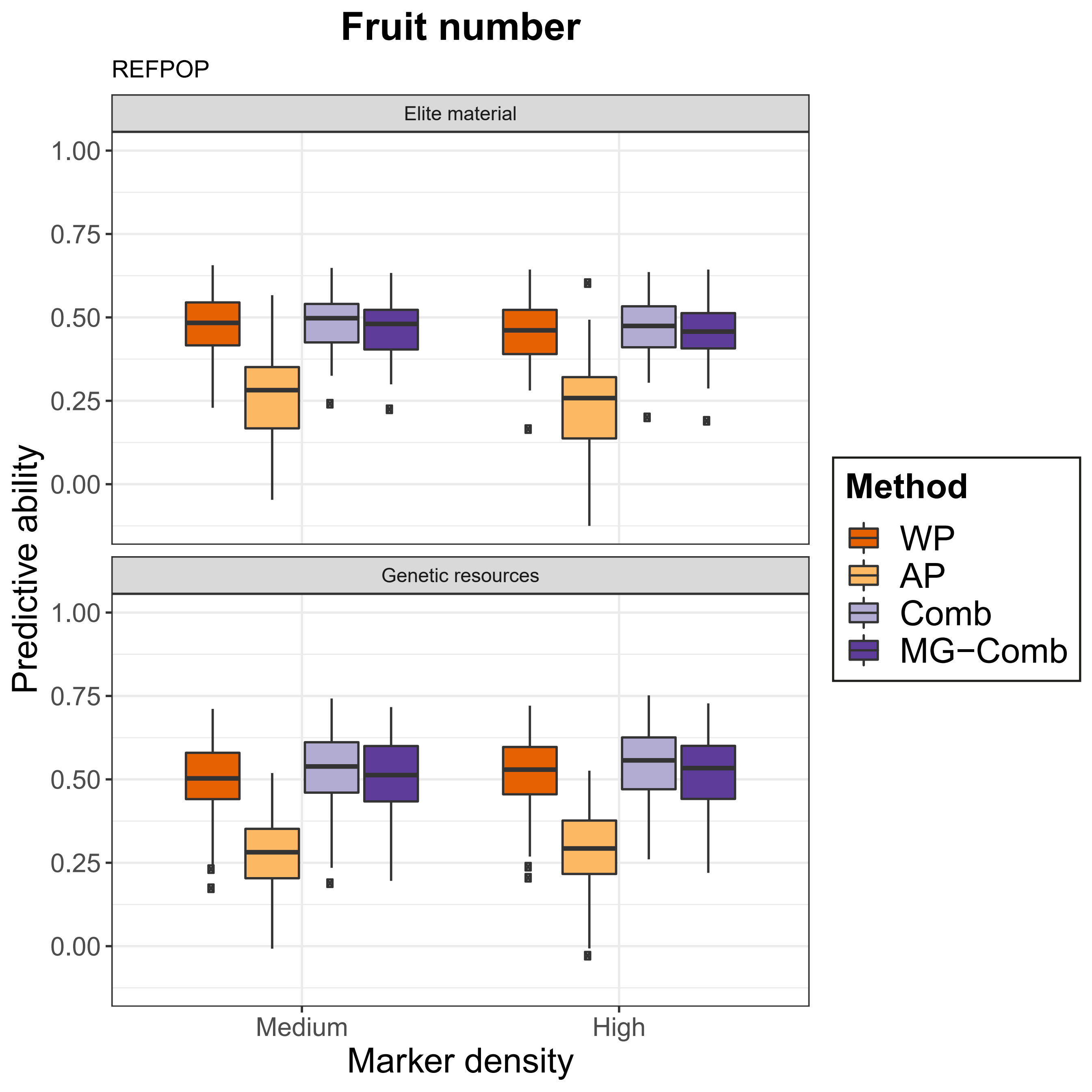

Fruit number
REFPOP
Elite material
Genetic resources
Predictive ability
1.00
0.75
0.50
0.25
0.00
Marker density
Medium
High
Method
WP
AP
Comb
MG−Comb

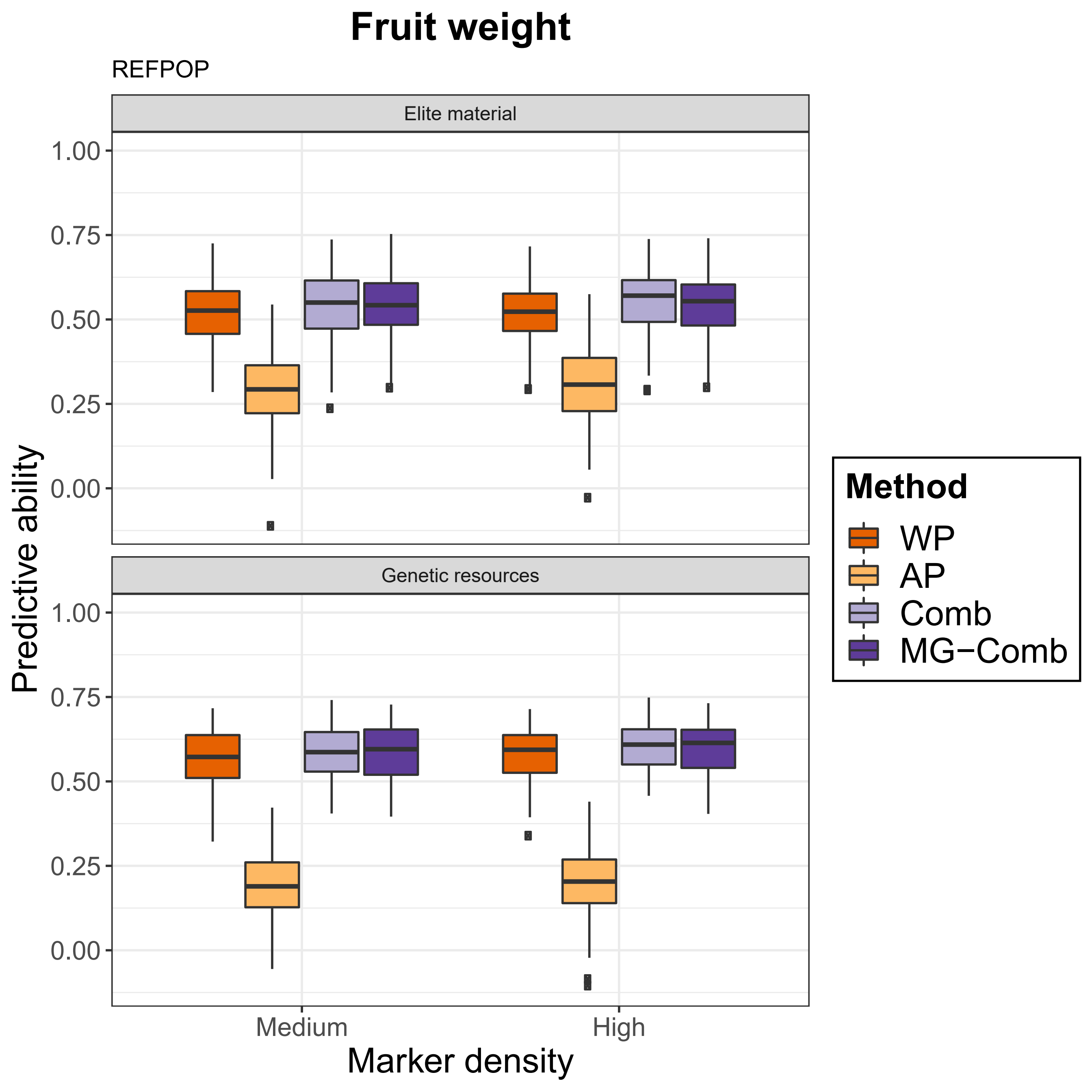

Fruit weight
REFPOP
Elite material
Genetic resources
1.00
0.75
0.50
0.25
0.00
Predictive ability
Medium
High
Marker density
Method
WP
AP
Comb
MG−Comb

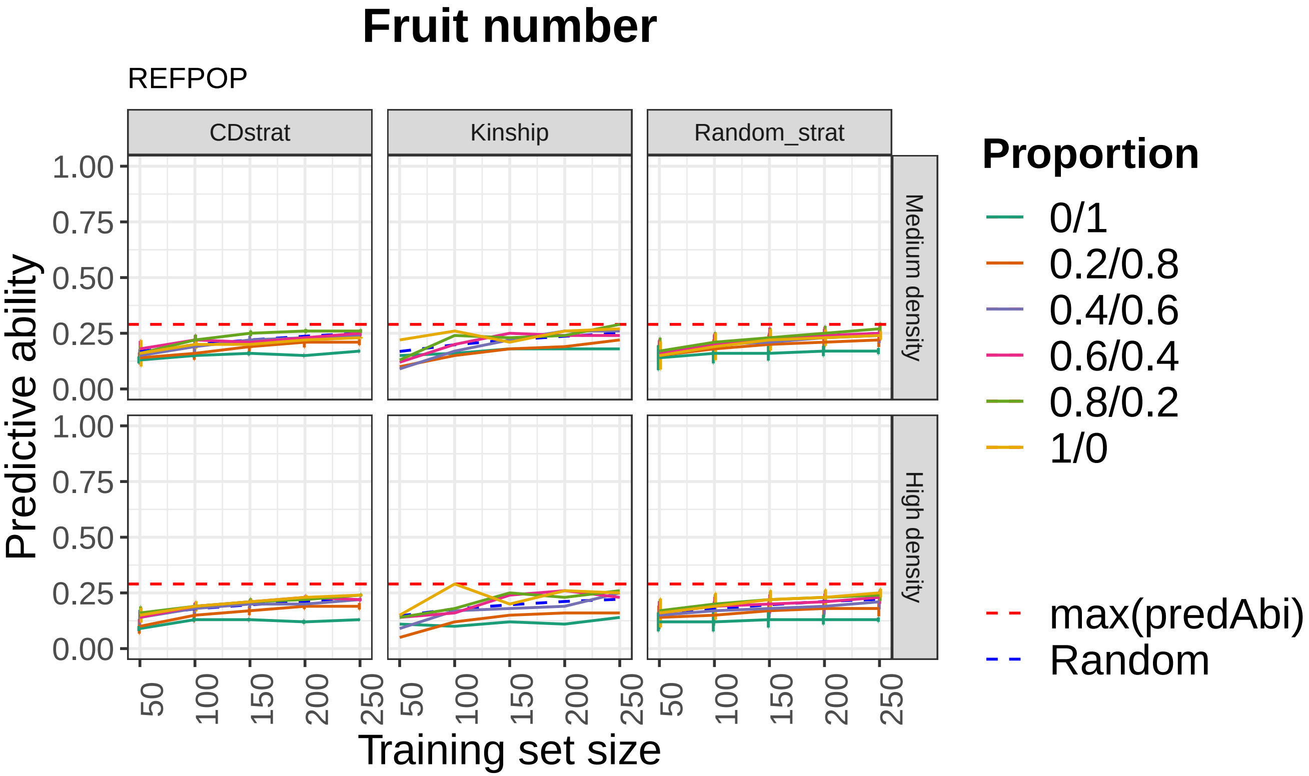

Fruit number
REFPOP
CDstrat
Kinship
Random_strat
Medium density
High density
Predictive ability
Training set size
1.00
0.75
0.50
0.25
0.00
50
100
150
200
250
Proportion
0/1
0.2/0.8
0.4/0.6
0.6/0.4
0.8/0.2
1/0
max(predAbi)
Random

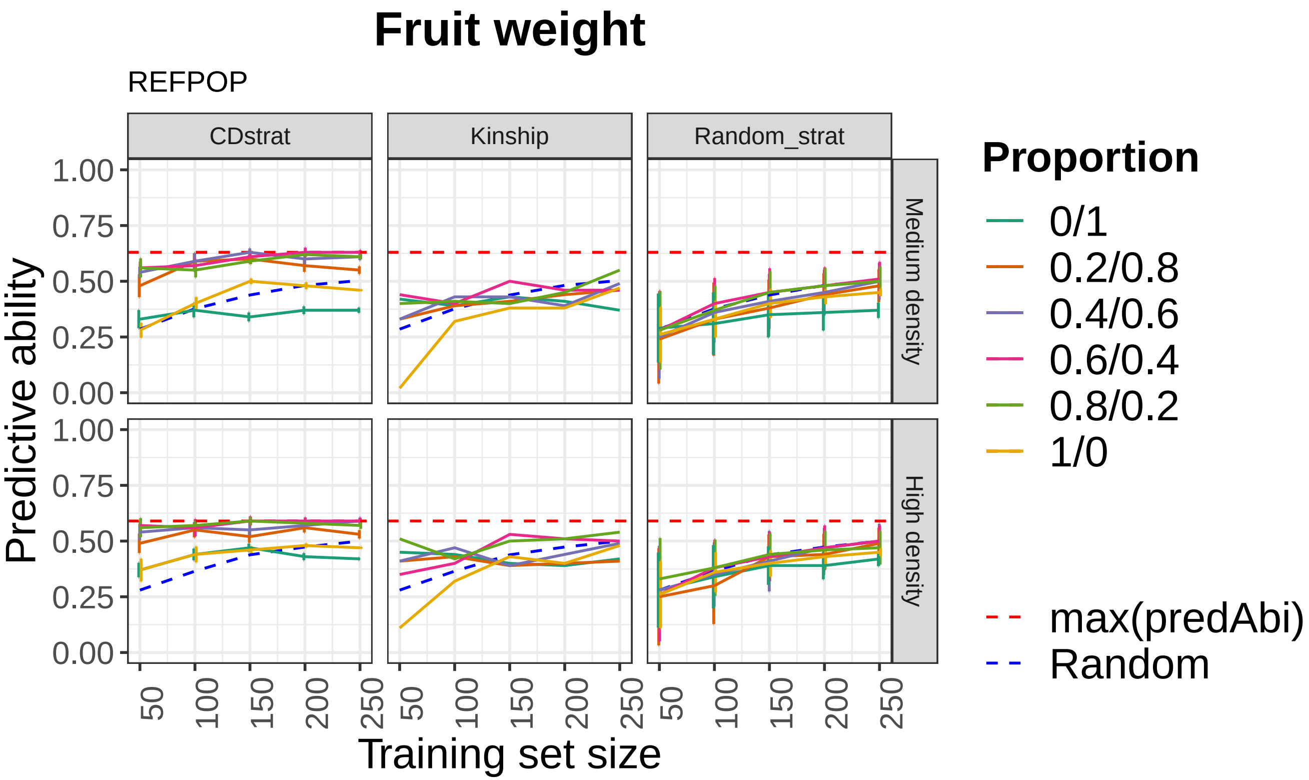

Fruit weight
REFPOP
CDstrat
Kinship
Random_strat
Medium density
High density
Predictive ability
1.00
0.75
0.50
0.25
0.00
50
100
150
200
250
Training set size
Proportion
0/1
0.2/0.8
0.4/0.6
0.6/0.4
0.8/0.2
1/0
max(predAbi)
Random
